# Contrasting patterns of genetic admixture explain the phylogeographic history of Iberian high mountain populations of midwife toads

**DOI:** 10.1101/2022.05.12.490531

**Authors:** Federica Lucati, Alexandre Miró, Jaime Bosch, Jenny Caner, Michael Joseph Jowers, Xavier Rivera, David Donaire-Barroso, Rui Rebelo, Marc Ventura

## Abstract

Multiple Quaternary glacial refugia in the Iberian Peninsula, commonly known as “refugia within refugia”, allowed diverging populations to come into contact and admix, potentially boosting substantial mito-nuclear discordances. In this study, we employ a comprehensive set of mitochondrial and nuclear markers to shed light onto the drivers of geographical differentiation in Iberian high mountain populations of the midwife toads *Alytes obstetricans* and *A. almogavarii* from the Pyrenees, Picos de Europa and Guadarrama Mountains. In the three analysed mountain regions, we detected evidence of extensive mito-nuclear discordances and/or admixture between taxa. Clustering analyses identified three major divergent lineages in the Pyrenees (corresponding to the eastern, central and central-western Pyrenees), which possibly recurrently expanded and admixed during the succession of glacial- interglacial periods that characterised the Late Pleistocene, and that currently follow a ring-shaped diversification pattern. On the other hand, populations from the Picos de Europa mountains (NW Iberian Peninsula) showed a mitochondrial affinity to central-western Pyrenean populations and a nuclear affinity to populations from the central Iberian Peninsula, suggesting a likely admixed origin for Picos de Europa populations. Finally, populations from the Guadarrama Mountain Range (central Iberian Peninsula) were depleted of genetic diversity, possibly as a consequence of a recent epidemic of chytridiomycosis. This work highlights the complex evolutionary history that shaped the current genetic composition of high mountain populations, and underscores the importance of using a multilocus approach to better infer the dynamics of population divergence.

## Introduction

Patterns of population structure and genetic divergence within species primarily result from their evolutionary history and contemporary dispersal capability, and these processes are in turn responsible of generating specific phylogeographic signatures [1]. The description of these signatures return valuable information on the mechanisms underlying spatial patterns of contemporary genetic diversity and structure, and this knowledge can ultimately help in the conservation and management of species and populations under global change [2].

Organisms with limited dispersal capacity and a typically allopatric type of speciation, such as many amphibians, generally present complex genetic signals at lineage borders resulting in reticulate patterns [3–5]. A number of mechanisms are involved in the generation of reticulate patterns in phylogeography, among them contact zones, hybridisation and introgression [6]. Such mechanisms, among others, may result in the formation of discordant patterns of variation among genetic markers (i.e. mito-nuclear discordances, e.g. [4, 7, 8]). As a consequence, these discordances represent a powerful source of insights into the evolutionary history of species.

Quaternary climatic fluctuations had a major impact on worldwide biotas and organisms, reshaping the distribution of species and modelling their genetic structure [9, 10]. In Europe, it is acknowledged that the Iberian Peninsula has served as one of the most important refugia for temperate species during periods of climatic instability [11]. Growing evidence has revealed that in this region, characterized by a great variety of climatic and ecological conditions, multiple isolated refugia were present, which promoted genetic diversification and ultimately determined complex phylogeographic patterns in a number of taxa (the “refugia within refugia” scenario; [11, 12]). In this context, Iberian mountain ranges played a major role in favouring survival throughout the Pleistocene. The Iberian Peninsula has several mountain ranges that permitted flexibility and survival of populations through altitudinal shifts, allowing for movements up or down the mountains in search of suitable microclimates as the temperatures changed [11, 13]. Nevertheless, there is a lack of knowledge on the phylogeographic history of European species with Iberian distribution to understand the role of glacial refugia throughout the different mountain regions within Iberia [11].

The common midwife toad (*Alytes obstetricans*) is a small anuran widely distributed in central and western Europe. It is a common species that inhabits a wide variety of habitats between sea level and 2 400 meters above sea level (m.a.s.l.) in the Pyrenees [14, 15]. Major threats to the species are related to habitat loss and fragmentation through pollution, commercial and agricultural development, introduction of non-native species, and more recently emergent diseases such as chytridiomycosis, which is caused by the chytrid fungus *Batrachochytrium dendrobatidis* (*Bd*) [14–16]. *Bd* has led to episodes of mass mortality in *A. obstetricans*, which is known to be highly susceptible to the pathogen [16–18]. Furthermore, fatal outbreaks of chytridiomycosis in Iberia were more frequent at higher elevations [19].

*Alytes obstetricans* represents an excellent biological system to study phylogeographic patterns derived from post-glacial expansion, such as contact zones and hybridisation. The species shows strong genetic subdivisions in the Iberian Peninsula, indicative of past population isolation [20]. Until recently, *A. obstetricans* was defined by six divergent and geographically structured mtDNA haplogroups (named A to F; [20, 21]), delineating as many genetic lineages that probably interbreed in zones of secondary contact (Fig 1). Each of the six mitochondrial lineages corresponded to a unique nuclear microsatellite clade, except for lineage B that harboured two distinct microsatellite clusters [22]. More recently, the subspecies *almogavarii* (mtDNA lineages E-F) was proposed as an incipient species (i.e. *A. almogavarii*; [23]) by RAD-sequencing analysis, and mtDNA lineage E described as a novel subspecies (*A. a. inigoi*) [24].

**Fig 1.**
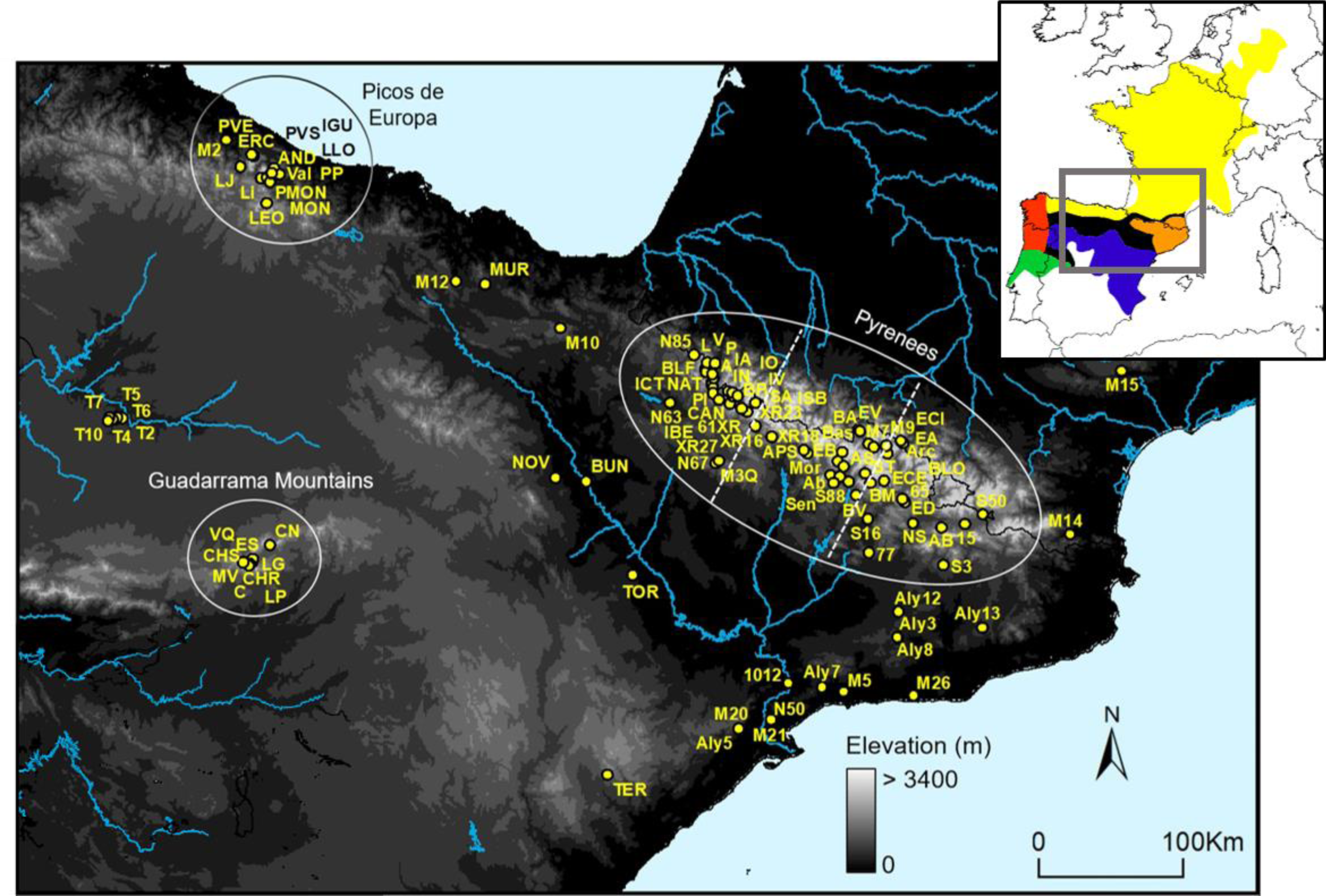
Geographic location of sampling sites for *Alytes obstetricans*/*almogavarii*. The three analysed mountain regions are circled in white. In the Pyrenees, dashed lines delimit the three geographic sections (eastern, central and western Pyrenees). For population codes and further information on sampling sites see S1 Table. The inset map shows the distribution of the main lineages: orange - ND4 haplogroups E-F (*A. almogavarii*), yellow - ND4 haplogroup B (*A. o. obstetricans*), blue - ND4 haplogroup A (*A. o. pertinax*), red - ND4 haplogroup C (*A. o. boscai*), green - ND4 haplogroup D (*A. o. boscai*), black - unclear (adapted from Dufresnes and Martínez- Solano [23]).

Previous studies revealed a complex scenario of relationships and admixture between lineages of *A. obstetricans*/*almogavarii*, evidencing the existence of contact and hybrid zones at some lineage borders [20, 23, 25, 26]. Yet, there has been little attempt on identifying the causes and consequences of such admixture.

Furthermore, none of these studies was specifically focused on high mountains, which are well-known hotspots of genetic diversity and have been shown to play a considerable role in separating well-differentiated intraspecific clades in numerous species (e.g. [4, 27]). Here, we employ a multilocus approach combined with comprehensive sample collection across four of the defined mtDNA genetic lineages to analyse the geographical differentiation in *A. obstetricans*/*almogavarii*, placing special focus on three high mountain regions in the Iberian Peninsula: Pyrenees, Picos de Europa and Guadarrama Mountain Range. Specifically, we aimed to (1) describe finer-scale geographical patterns of genetic diversity and structure in high mountain populations of *A. obstetricans*/*almogavarii*, compare different scenarios of population divergence and explore the role of glacial refugia; (2) identify putative contact zones and assess patterns of admixture; and (3) better delineate the geographic distribution of major genetic lineages. We also provide deeper analyses on the genetic status of populations that have been hit by a recent chytridiomycosis outbreak. To this end, we combined DNA sequence (mitochondrial and nuclear) and microsatellite data for the first time in an *A. obstetricans*/*almogavarii* study. Thus, herein we provide new insights on the recent evolutionary history and the present processes that have shaped and are currently shaping Iberian populations of *A. obstetricans*/*almogavarii*.

## Materials and Methods

### Sampling and DNA extraction

Sampling was conducted mainly in the period 2009–2018 across northern and central Iberian Peninsula, as it is known to host most of the genetic diversity of the species [20, 22]. We paid special emphasis on three mountain regions: Pyrenees, Picos de Europa mountains (Cantabrian Mountains, NW Iberian Peninsula) and Guadarrama Mountain Range (central Iberian Peninsula), although we also took some additional samples from neighbouring lowland localities (Fig 1, Tables 1 and S1). We sampled 51 sites in the Pyrenees (1010–2447 m.a.s.l.), 13 in Picos de Europa (1120–2079 m.a.s.l.), nine in Guadarrama Mountains (1527–1980 m.a.s.l.), and 32 sites in lowland areas (52–992 m.a.s.l.), totalling 890 individuals from 105 sampling sites (Table 1). Lowland populations were grouped according to their geographic proximity to the analysed mountain regions (Table 1). A subset of samples was used in a previous study [18].

**Table 1.** Genetic diversity parameters as estimated from microsatellites for each analysed mountain region and corresponding lowland areas.

|  | Alt. | N<sub>pops</sub> | N | Na | Ar | PAAr | H<sub>O</sub> | H<sub>E</sub> | N<sub>e</sub> | ND4 haps |
| --- | --- | --- | --- | --- | --- | --- | --- | --- | --- | --- |
| Eastern Pyrenees | 1010-2406 | 17 | 144 | 13.47 | 5.310 | 0.780 | 0.480 | 0.701 | 14-51 | 28(F) |
| Eastern Pyrenees – lowland | 62-948 | 9 | 21 | 8.588 | 5.310 | 0.830 | 0.460 | 0.691 | - | 11(F) |
| Central Pyrenees | 1211-2003 | 9 | 52 | 10.82 | 5.130 | 0.920 | 0.401 | 0.651 | 68 | 15(E), 4(B) |
| Central Pyrenees – lowland | 567 | 1 | 2 | - | - | - | - | - | - | 2(E) |
| Central-western Pyrenees | 1043-2447 | 25 | 284 | 14.12 | 5.240 | 0.990 | 0.377 | 0.708 | 12-84 | 61(B) |
| Central-western Pyrenees – lowland | 992 | 1 | 1 | - | - | - | - | - | - | 1(B) |
| Picos de Europa | 1120-2079 | 13 | 202 | 12.71 | 5.740 | 1.080 | 0.595 | 0.789 | 19-160 | 31(B) |
| Picos de Europa – lowland | 464-928 | 3 | 3 | - | - | - | - | - | - | 3(B) |
| Guadarrama | 1527-1980 | 9 | 112 | 7.529 | 3.940 | 0.580 | 0.442 | 0.613 | 5-15 | 25(A) |
| Guadarrama – lowland | 52-830 | 18 | 57 | 14.53 | 6.580 | 1.080 | 0.620 | 0.805 | 70 | 36(A), 2(B) |

Alt. – altitudinal range, N<sub>pops</sub> – number of populations, N – sample size for microsatellites, Na – mean number of alleles, Ar – allelic richness standardized for sample size, PAAr – rarefied private allelic richness standardized for sample size, H<sub>O</sub> – observed heterozygosity, H<sub>E</sub> – expected heterozygosity, N<sub>e</sub> – range of effective population size, ND4 haps – occurrence and code (in parentheses) of mitochondrial ND4 haplogroups identified in each deme.

Tissue samples were collected via tail clipping in the case of larvae and toe clipping in the case of adults. Samples were stored in absolute ethanol and maintained at -20 °C. Genomic DNA was extracted using QIAGEN DNeasy Blood & Tissue Kit (Qiagen, Hilden, Germany) according to the manufacturer’s protocol, or following the HotSHOT method [28], in a final volume of 100 µl for toe clips and 250 µl for tail clips.

### Nuclear and mitochondrial genes sequencing

We amplified five gene regions, including four mitochondrial fragments (cytochrome *b* gene – cyt-*b*; 12S rRNA gene – 12S; 16S rRNA gene – 16S; and NADH dehydrogenase subunit 4 gene and adjacent tRNAs – ND4) and one nuclear gene (β- fibrinogen intron 7 – β-fibint7). Details on primers used and amplification conditions are provided in S1 Appendix.

Resulting sequences were aligned using the ClustalW algorithm in MEGA 7 with default settings [29]. Phasing of the nuclear intron β-fibint7 was performed following Gonçalves et al. [20]. Briefly, we used the program SeqPHASE [30] to format the input files, and the Bayesian algorithm implemented in PHASE 2.1.1 [31, 32] to infer phased haplotypes. PHASE was run three times using default values, to check for consistency of haplotype estimation across runs.

### Microsatellite screening and estimation of genetic diversity

878 individuals from 102 localities were screened for 17 previously characterised microsatellite loci combined in five multiplexes [21]. Fragments were sized with LIZ-500 size standard and binned using Geneious 11.0.5 [33].

The presence of potential scoring errors, large allele dropout and null alleles was tested using MICRO-CHECKER 2.2.3 [34]. We tested for linkage disequilibrium between loci and for departures from Hardy-Weinberg equilibrium (HWE) in each population and for each locus using GENEPOP 4.2 [35]. The Bonferroni correction was applied to adjust for multiple comparisons (α = 0.05; [36]).

Genetic diversity parameters were calculated for populations with ≥ five genotyped individuals and for each analysed mountain region and corresponding lowland areas. Observed (HO) and expected heterozygosity (HE) and mean number of alleles (Na) were obtained with GenAlEx 6.5 [37]. Allelic richness (Ar) standardized for sample size and rarefied private allelic richness (PAAr – calculated only at the genetic cluster level) were calculated in HP-RARE 1.1 [38]. We estimated the inbreeding coefficient (FIS) within each population with GENETIX 4.05.2 [39].

### Phylogenetic analyses

For ND4, we estimated genealogic relationships among haplotypes using Haploviewer [40]. The optimal nucleotide-substitution model was determined by jModelTest 2.1.3 [41], under the Akaike Information Criterion (AIC). The phylogeny was estimated with RAxML 7.7.1 [42] using a Maximum Likelihood (ML) approach. The program was run with a gamma model of rate heterogeneity and no invariant sites (GTRGAMMA), applying 1 000 bootstrap replicates. The best tree was selected for haplotype network construction in Haploviewer, based on all sequences retrieved from GenBank and this study. Furthermore, genetic diversity parameters, namely number of haplotypes (H) and polymorphic sites (S), as well as haplotype (Hd) and nucleotide (Π) diversity indices, were estimated for the whole dataset and for each analysed mountain region and corresponding lowland areas using DnaSP 6.11.01 [43].

In order to describe the affinities of the different genetic lineages described in *A. obstetricans*/*almogavarii*, we constructed a species tree with the five partially sequenced genes using the multispecies coalescent approach in *BEAST, as implemented in BEAST 2.6.2 (S1 Fig) [44]. The substitution model for each marker was determined by jModelTest. We set a strict molecular clock model and a Yule speciation prior. The clock and tree models for mtDNA markers were linked. We defined five groups in *A. obstetricans*/*almogavarii*, corresponding to the four main population lineages identified in the haplotype network (A, B, E, F) and further subdividing lineage B into two groups (central-western Pyrenees and Picos de Europa; see Results). Since *BEAST does not take into account the possibility of gene flow between lineages, samples from localities where more than one mtDNA haplogroup was found were excluded from the analysis [20]. Samples of *A. maurus* were used as outgroup. Two independent MCMC chains were run for 200 million generations each, sampling trees and parameter estimates every 5 000. Tracer 1.6 [45] was used to check for stationarity and convergence of MCMC chains. A maximum clade credibility tree was constructed in TreeAnnotator 2.6.2 and visualized using FigTree 1.4.3 (http://tree.bio.ed.ac.uk/software/figtree), discarding the first 10% generations as burn-in, while the node heights were set to mean heights.

### Genetic structure

For microsatellites, population structure was inferred using different approaches (S1 Fig): a) a Discriminant Analysis of Principal Components (DAPC) and a Principal Component Analysis (PCA), using the ADEGENET package 2.1.1 [46] in R 3.5.1 [47]; b) a Bayesian cluster analysis implemented in STRUCTURE 2.3.4 [48]; c) a neighbour-joining (NJ) tree using the program POPTREEW [49]; and d) a spatial-based clustering approach implemented in the R package Tess3R 1.1.0 [50], which incorporates geographic coordinates in estimating sample ancestry coefficients. Details on models and parameters used are outlined in S1 Appendix.

To further explore the genetic differentiation between the defined genetic units, we calculated pairwise FST for both ND4 and microsatellites. Computations were performed in Arlequin 3.5.2.2 [51] in the case of ND4 and in GenAlEx in the case of microsatellites, with 1 000 permutations to assess statistical significance. Similarly, to partition genetic variability at different hierarchical levels (among genetic clades, among populations within clades and within populations), we conducted an analysis of molecular variance (AMOVA) as implemented in Arlequin, using 10 000 permutations to assess significance of variance components [52]. To better understand the spatial pattern of population differentiation detected in the Pyrenees (see Results), we correlated genetic distances (FST) of Pyrenean populations either with the Euclidean distance among populations, and with the geographic distance suggested by PCA analysis, which followed a ring-like pattern around the Pyrenean chain assuming no direct gene flow across the central Pyrenean axis (see e.g. [53] for a similar approach). The analysis was performed in GenAlEx with 1 000 permutations, including only populations with ≥ five genotyped individuals (N = 29).

### Effective population size

The sibship assignment method implemented in Colony 2.0.6.5 was used to calculate the effective population size (Ne) of populations with ≥ 15 genotyped individuals (S1 Fig) [54]. This software uses a maximum likelihood method to conduct parentage and sibship inference to estimate Ne and can accommodate null alleles and other genotyping errors. The program was run with the parameters specified in Lucati et al. [55]. During sampling care was taken to minimize the effects of sampling individuals belonging to the same clutch, by collecting samples from several spots within the same sampling site.

### Demographic history

The approximate Bayesian computation (ABC) approach, as implemented in the software DIYABC 2.1.0 [56], was used to reconstruct the history of divergence among the genetic lineages identified in *A. obstetricans*/*almogavarii*. Only high mountain populations were included in the analysis, as we were interested in understanding the evolutionary history of the three high mountain areas described above (S1 Fig). We grouped populations into five groups according to their geographic location. We also incorporated the information from previous studies dealing with the phylogenetics and population structure of the different subspecies, which point to a shared origin between populations from the central and eastern Pyrenees (mitochondrial ND4 haplogroups E and F in Gonçalves et al. [20] and microsatellite clusters F1n and F2n in Maia-Carvalho et al. [22]): populations from the Picos de Europa mountains (PEU) and the central-western Pyrenees (CWPY; corresponding to subspecies *A. o. obstetricans*) formed two separate groups, whereas populations from the central Pyrenees (CPY), eastern Pyrenees (EPY; corresponding to the putative incipient species *A. almogavarii*) and Guadarrama Mountains (GUA; corresponding to subspecies *A. o. pertinax*) were subdivided into three groups. We generated five different scenarios (Fig 2a): (1) null model with all groups diverging simultaneously from a common ancestor, except for lineage CPY that originates from EPY; (2) sequential splitting model directly following results from STRUCTURE and DAPC analyses, where on the one hand populations from the Pyrenees, and on the other GUA and PEU populations share a common origin; (3) similar to the second one, but predicts a common origin of PEU and CWPY populations, as suggested by ND4 haplotype network; (4) same as the previous scenario, with a common origin between populations belonging to ND4 haplogroups A and B, as suggested by *BEAST analysis; and (5) where the PEU lineage was created by admixture of CWPY and GUA populations. We performed the computations both combining microsatellites and ND4 data, and separately for microsatellites. Further details on model specifications and run parameters are outlined in S1 Appendix and S2 Table.

**Fig 2.**
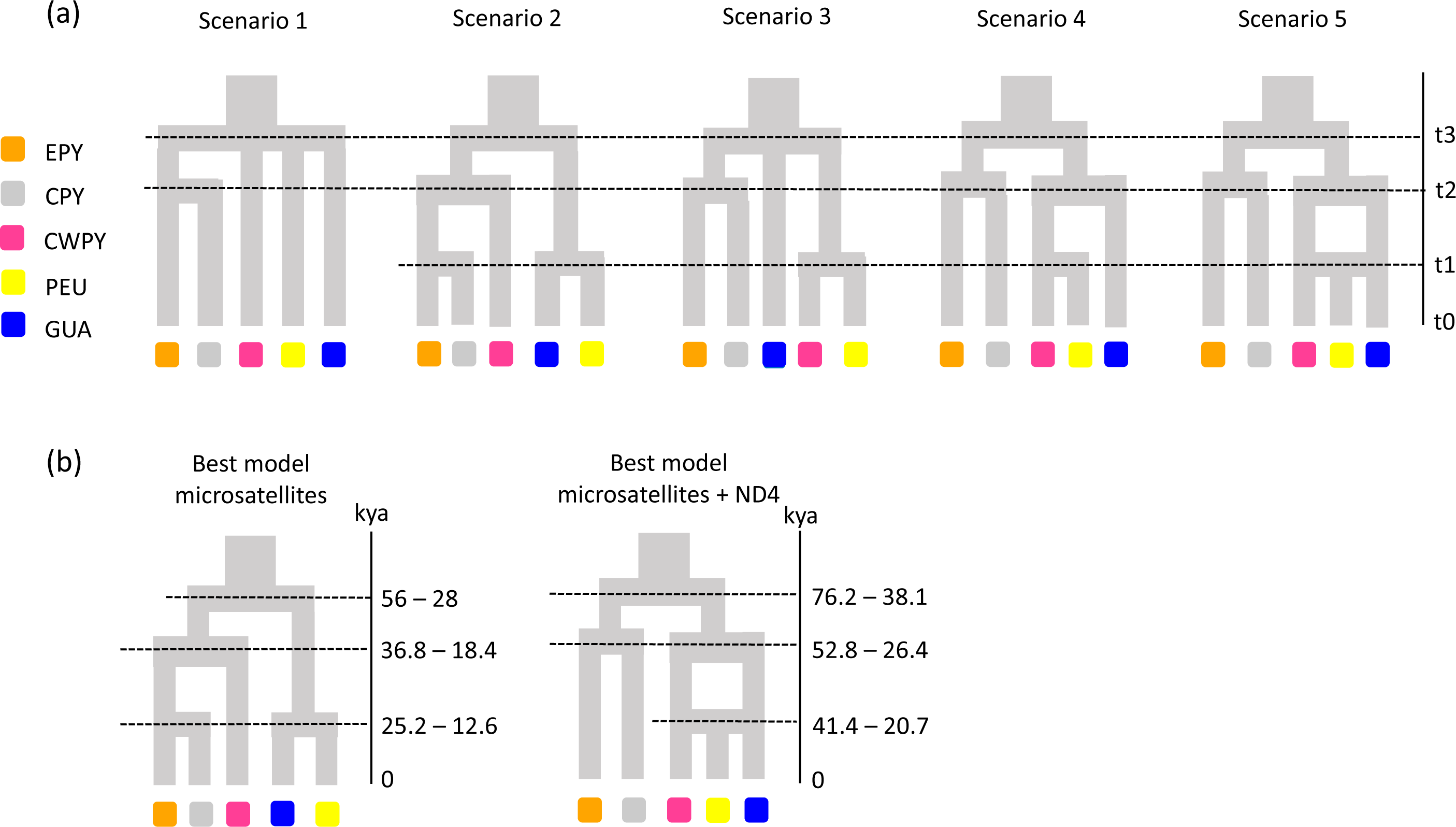
Phylogeographic scenarios tested in DIYABC and best supported models. (a) Scenarios tested in DIYABC considering the three analysed high mountain areas. (b) The best supported scenarios, namely number 2 when considering only microsatellites and number 5 when considering both mtDNA (ND4) and microsatellite markers, with the estimated divergence times (t1 – t3) of each split. Population groups were defined on the basis of their geographic distribution and the results from clustering analyses: eastern Pyrenees (EPY, orange), central Pyrenees (CPY, grey), central-western Pyrenees (CWPY, pink), Picos de Europa mountains (PEU, yellow), and Guadarrama Mountain Range (GUA, blue).

## Results

### Sequence variation and genetic diversity

The nuclear DNA alignment (β-fibint7) included 20 sequences of 605 bp, while for the mitochondrial dataset we obtained 219 sequences of 654 bp for ND4, 40 sequences of 325 bp for cyt-*b*, 42 sequences of 341 bp for 12S, and 40 sequences of 579 bp for 16S (S1 Table). With regard to ND4, overall haplotype (Hd) and nucleotide (Π) diversities were 0.930 ± 0.008 and 0.020 ± 0.0008, respectively. Across the analysed mountain regions, Guadarrama Mountains showed lower ND4 genetic diversity values (Hd = 0.290 ± 0.109, Π = 0.001 ± 0.0002) compared to the other regions (eastern Pyrenees: Hd = 0.770 ± 0.039, Π = 0.002 ± 0.0003; central Pyrenees: Hd = 0.724 ± 0.101, Π = 0.014 ± 0.003; central-western Pyrenees: Hd = 0.805 ± 0.027, Π = 0.002 ± 0.0002; Picos de Europa: Hd = 0.602 ± 0.091, Π = 0.003 ± 0.001).

For microsatellite loci, we did not find evidence of large allele dropout, stuttering or null allele artefacts. Similarly, no significant linkage disequilibrium or departures from HWE across populations and loci were detected after applying the Bonferroni correction. Observed (HO) and expected heterozygosity (HE) ranged from 0.197 to 0.794 and from 0.222 to 0.720, respectively (mean HO = 0.473, mean HE = 0.530; S1 Table). Mean number of alleles (Na) and allelic richness (Ar) varied from 2.059 to 7.824 and from 1.270 to 5.620, respectively (mean Na = 4.440, mean Ar = 1.873), and the inbreeding coefficient (FIS) ranged from -0.110 to 0.403 (mean = 0.157). Among the analysed mountain regions, Picos de Europa and the eastern Pyrenees were the richest regions in terms of genetic diversity, whereas Guadarrama Mountains was the poorest, with the central and central-western Pyrenees presenting intermediate values (Table 1). With regard to lowland areas, populations located in the Duero and Ebro basins (herein referred to as Guadarrama lowland localities) were the most diverse, while eastern Pyrenean localities were the least diverse.

### Phylogenetic analyses

From the 219 individuals analysed for the ND4 gene we identified 34 haplotypes, of which 22 are newly described (Fig 3, S1 Table). Haplotypes were defined by 61 polymorphic sites, of which 49 were parsimony informative. The majority of newly described haplotypes were found in the Pyrenees (13), whereas only three and one haplotypes were detected in Picos de Europa and Guadarrama Mountains, respectively; the remaining five haplotypes were found in lowland localities. The haplotype network showed that haplotypes clustered into four well- differentiated haplogroups with a strong association with geography (Figs 3 and S2a): haplogroup A included sequences from Guadarrama and corresponding lowland areas (corresponding to populations of *A. o. pertinax*), haplogroup F included sequences from the eastern Pyrenees (corresponding to populations of *A. almogavarii*), haplogroup E corresponded to populations from the central Pyrenees (*A. a. inigoi*), and haplogroup B (corresponding to populations of *A. o. obstetricans*) included sequences from the central-western Pyrenees and Picos de Europa. Within haplogroup B, sequences corresponding to central-western Pyrenean populations were clearly separated from those from Picos de Europa. In addition, in four localities we detected the presence of more than one haplogroup (Figs 3 and S2a, S1 Table): haplogroups A and B were found to co-occur in population Fte. Nueva de Bardales (T2), whereas both haplogroups B and E were found in populations Plano de Igüer (PI), Balsa Pertacua (BP) and Ibón de los Asnos (61XR).

**Fig 3.**
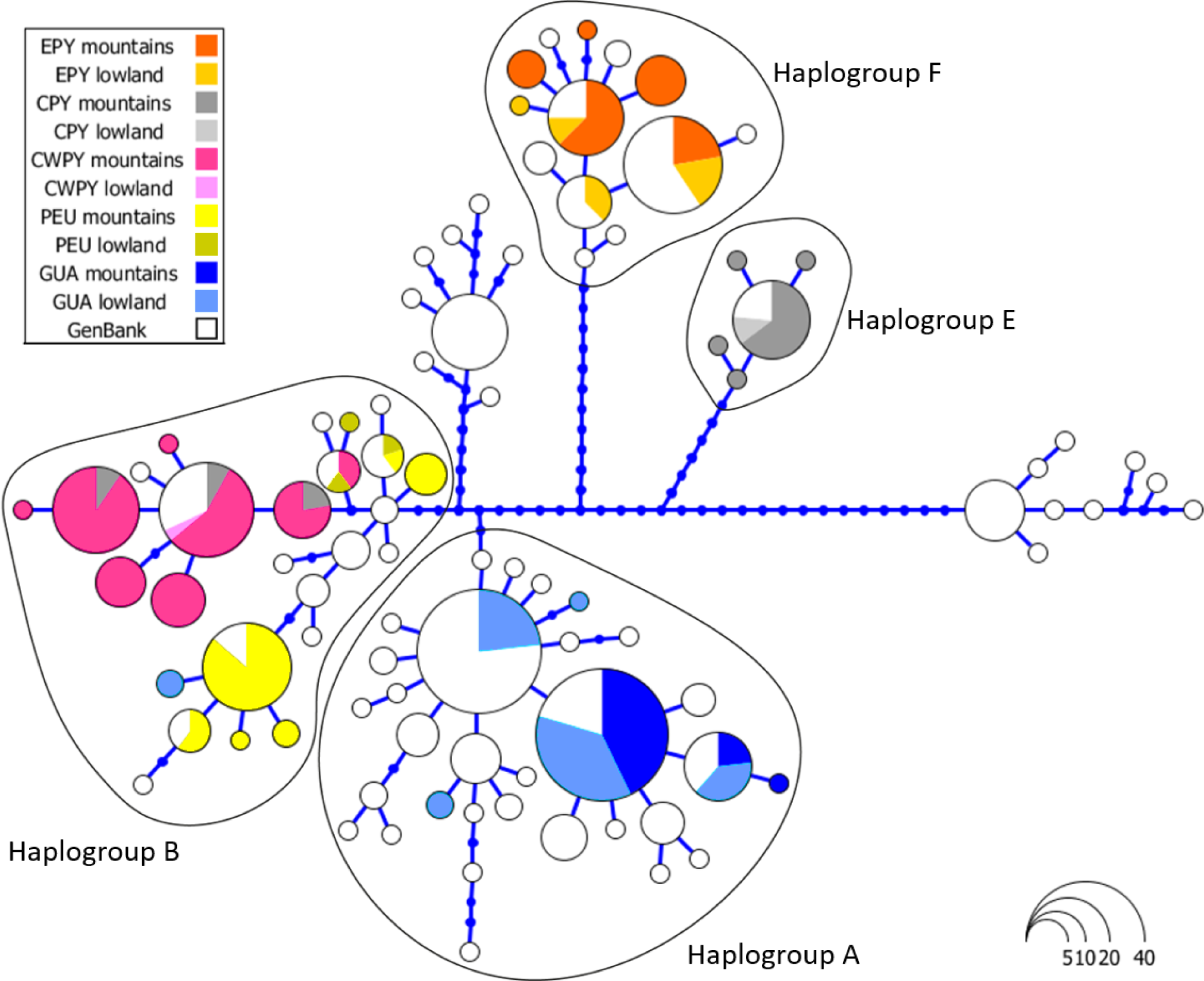
Haplotype network of ND4 sequences analysed in *Alytes obstetricans*/*almogavarii*. Each circle represents a unique haplotype and the circle area is proportional to the number of sequences of a given haplotype. Blue dots correspond to inferred unsampled haplotypes. Sequences depicted in white were retrieved from GenBank. The analysed high mountain regions and corresponding lowland areas are indicated by different colours: eastern Pyrenees (EPY, orange), central Pyrenees (CPY, grey), central-western Pyrenees (CWPY, pink), Picos de Europa mountains (PEU, yellow), and Guadarrama Mountain Range and corresponding lowland areas (GUA, blue).

In order to describe the affinities between the different haplogroups, we built a consensus tree with the five sequenced gene fragments using a multispecies coalescent approach (species tree; *BEAST). The tree placed lineage E at the base of the *A. obstetricans*/*almogavarii* lineages, with lineage A as sister to lineage B. Finally, lineage B split into central-western Pyrenees and Picos de Europa groups (S3 Fig).

### Genetic structure and effective population sizes

All the clustering analyses performed on microsatellites recovered congruent results (Figs 4, 5 and S2). Regarding DAPC analysis, the optimum number of clusters was inferred to be 7 (Figs 4c, S2b and S4b). The inferred clusters were consistent with the geographic location of populations. A closer look from K = 2 to K = 7 revealed a spatial hierarchical genetic structure (S4a Fig), with a first split between the central- western Pyrenees and all other populations, and a subsequent subdivision of this second group into central Pyrenees–eastern Pyrenees and Guadarrama and corresponding lowland populations–Picos de Europa. At K = 4, populations from the Guadarrama Mountain Range split from lowland populations located in the Duero and Ebro basins and Picos de Europa Mountains, whereas at K = 5 there was a split between the central and eastern Pyrenees. The following splits occurred within the central-western Pyrenean group. According with DAPC analysis, the PCA grouped populations according to their geographic distribution rather than by described subspecies (Fig 4d). PC1 separated the Pyrenees from the Picos de Europa and Guadarrama, whereas PC2 distributed Pyrenean populations along the Pyrenean chain and contributed to the separation of the Picos de Europa and Guadarrama. STRUCTURE analysis showed patterns of genetic differentiation similar to those inferred by PCA, and identified K = 2 as the best clustering solution (Figs 4a and S5): the first cluster grouped together all Pyrenean populations and the second cluster included peninsular ones. A smaller peak was detected at K = 7 (Fig 4a, 5 and S5b), in agreement with DAPC-inferred clusters, but segregating populations located in the Duero and Ebro basins from Guadarrama Mountains, while splitting central-western Pyrenean populations into two groups. This partitioning into seven clusters was also supported by results of Tess3R analyses (S6 Fig). Furthermore, a higher level of genetic admixture was detected in lowland populations than in highland populations (Figs 5 and S6). More into detail, lowland *A. o. pertinax* populations generally presented the highest degree of admixture, whereas in the Pyrenees moderate levels of admixture were detected between *A. almogavarii* lineage E and *A. obstetricans* in the west, as well as between the two *A. obstetricans* lineages, but not between the two *A. almogavarii* lineages (Fig 5). In order to describe the segregation history amongst the seven groups identified by STRUCTURE, we built a NJ tree based on net nucleotide distances: the tree started at K = 3 with the separation of Guadarrama and corresponding lowland populations from Picos de Europa and the Pyrenees, then at K = 4 populations from the central and eastern Pyrenees split from central-western Pyrenean populations, whereas the following splits occurred within the different major groups (Fig 4b). We also built a second NJ tree inferred from microsatellite- based DA distances over all populations that identified five discrete lineages, which match with ND4-inferred lineages, except that populations included in haplogroup B formed two separate groups (S2d and S7 Figs). Nevertheless, the tree suggested that the pairs of lineages central Pyrenees–eastern Pyrenees and Guadarrama–Picos de Europa were more closely related with each other than with the lineage of central-western Pyrenees, which in turn appeared more distant from the rest of the lineages.

**Fig 4.**
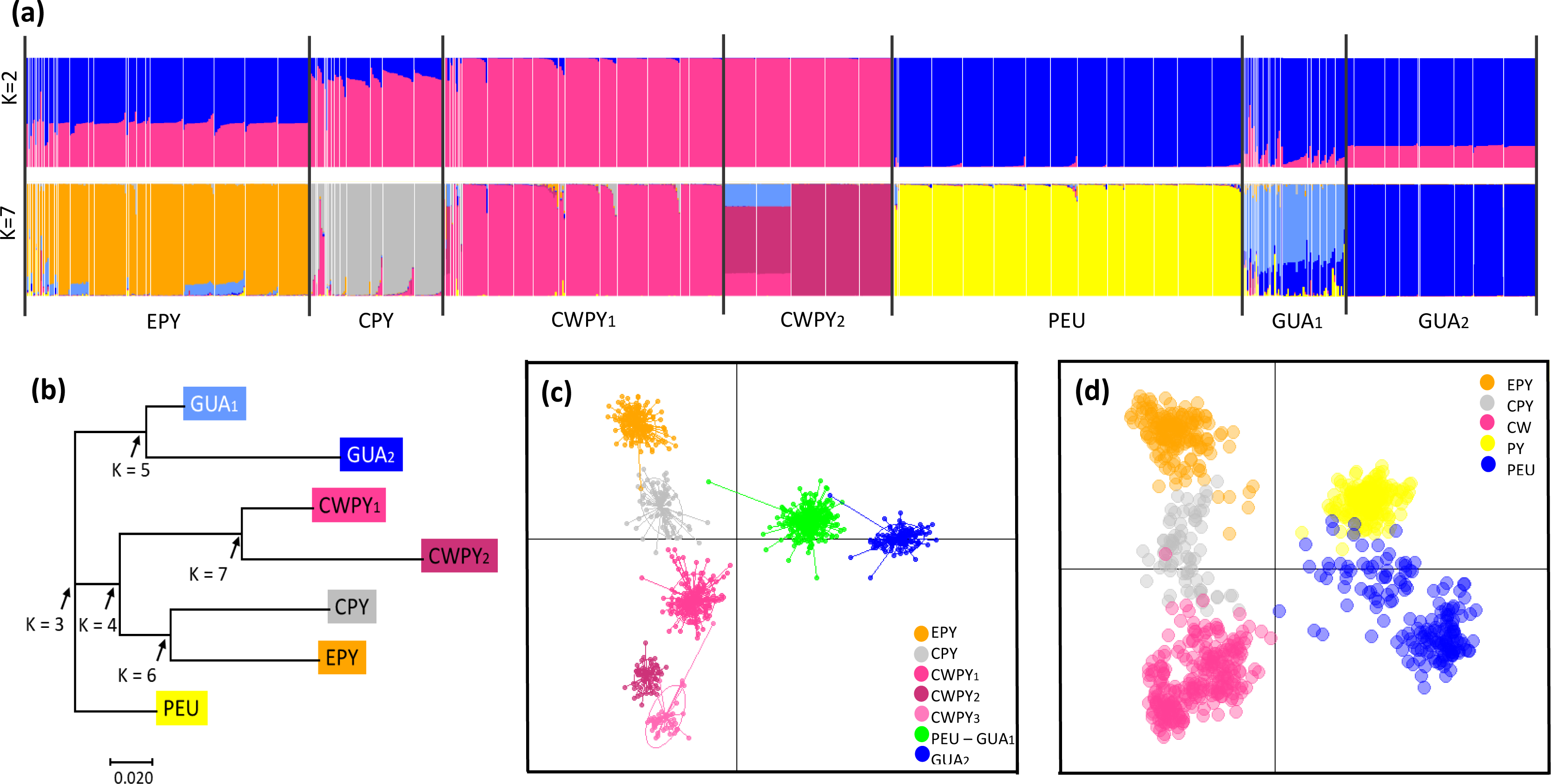
Results of clustering analyses of *Alytes obstetricans*/*almogavarii* populations based on microsatellite data. (a) STRUCTURE barplots of membership assignment for K = 2 and K = 7. Each individual is represented by a vertical bar corresponding to the assignment probabilities to the K cluster. White lines separate populations and black lines separate clusters. (b) Neighbour-joining tree based on net nucleotide distances among the seven clusters inferred by STRUCTURE. Arrows indicate the sequence of differentiation when K increases. (c) Summary plot from Discriminant Analysis of Principal Components (DAPC) for K = 7 genetic clusters. Dots represent individuals and genetic clusters are shown as inertia ellipses. (d) Plot obtained in the Principal Component Analysis (PCA). Each dot represents one individual. Labels indicate the different genetic clusters: eastern Pyrenees (EPY, orange), central Pyrenees (CPY, grey), central-western Pyrenees (CWPY, pink), Picos de Europa mountains (PEU, yellow), and Guadarrama Mountain Range (GUA, blue).

**Fig 5.**
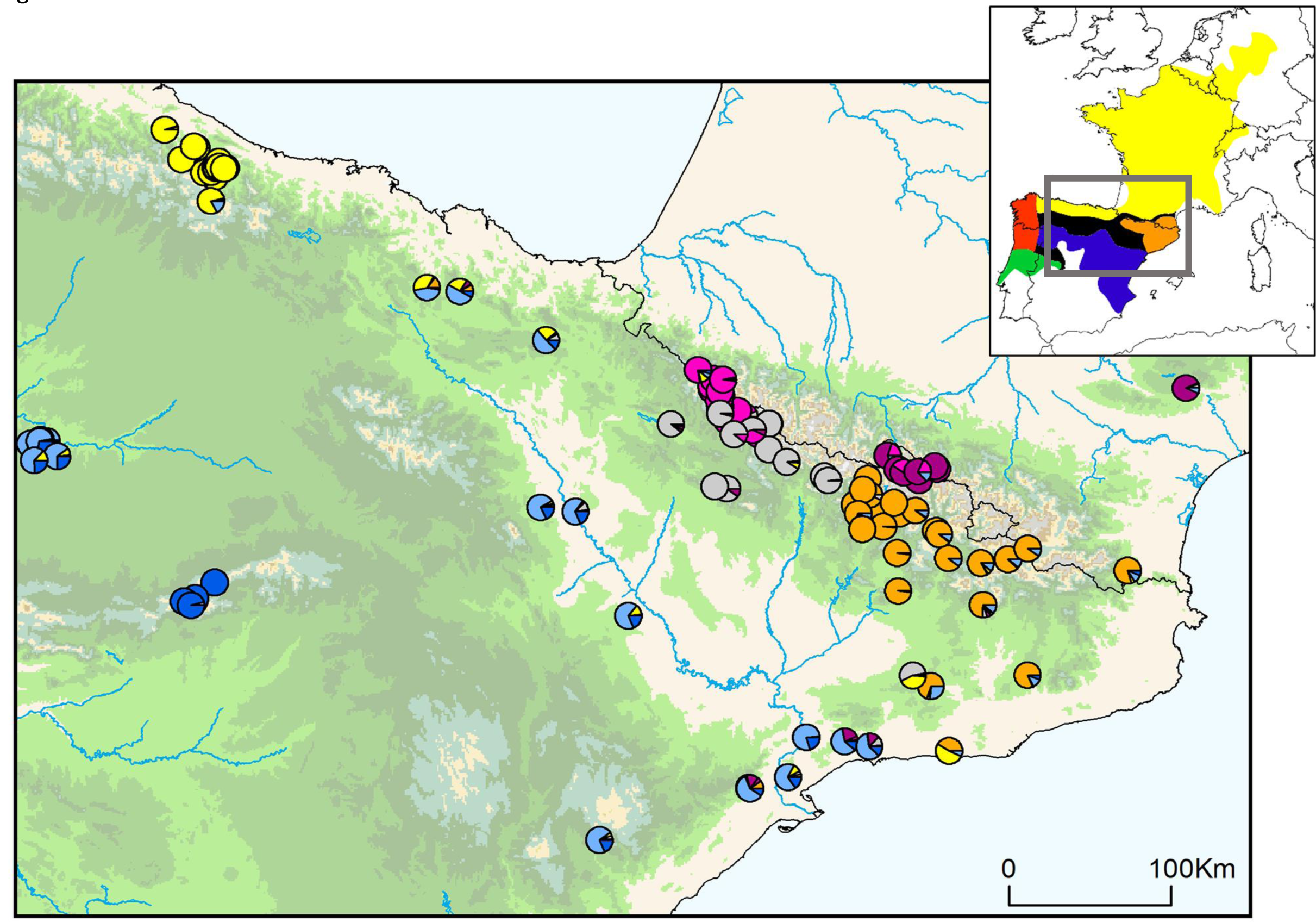
Results of Bayesian clustering analysis (STRUCTURE) for K = 7 microsatellite groups for *Alytes obstetricans*/*almogavarii*. Sampled populations are represented by pie charts highlighting the population cluster membership obtained in STRUCTURE. The inset map shows the distribution of the main lineages: orange - ND4 haplogroups E-F (*A. almogavarii*), yellow - ND4 haplogroup B (*A. o. obstetricans*), blue - ND4 haplogroup A (*A. o. pertinax*), red - ND4 haplogroup C (*A. o. boscai*), green - ND4 haplogroup D (*A. o. boscai*), black - unclear (adapted from Dufresnes and Martínez- Solano [23]).

The test of significance of pairwise FST values between the seven lineages defined by STRUCTURE was significantly different from zero (P < 0.01; S3 Table), indicating significant genetic differences. ND4-based pairwise FST values ranged from 0.951 for eastern Pyrenees–Guadarrama Mountains to 0.089 for Guadarrama Mountains–Guadarrama lowlands, whereas microsatellite-based pairwise FST values ranged from 0.222 for Guadarrama Mountains–central-western Pyrenees to 0.060 for Guadarrama lowlands–Picos de Europa. Accordingly, AMOVA analyses suggested significant structure among the seven genetic lineages (P < 0.001; S4 Table). The proportion of variation attributable to differences among lineages was lower in microsatellites (26.171%) than in ND4 (84.945%), possibly as a result of gene flow and the inability of mtDNA to detect it. Conversely, variation among individuals within populations was low for ND4 (8.736%) and high for microsatellite (56.425%) markers, as expected for polymorphic loci such as microsatellites.

In the Pyrenees, microsatellite genetic differentiation followed a ring-like pattern, being maximal between lineages EPY (orange) and CWPY (pink; at opposite extremities on PCA axis 2, Fig 4c-d), even though they are geographically proximate in the central and eastern Pyrenees. To evaluate this hypothesis of ring-shaped isolation by distance, we correlated genetic distances (FST obtained from microsatellites) of Pyrenean populations to different types of geographic distance. We obtained a weak isolation by distance pattern between genetic differentiation and Euclidean geographic distance (Mantel’s R = 0.185, P = 0.002; Fig 6). In contrast, we detected a strong correlation between genetic differentiation and the geographic distance suggested by PCA analysis (Fig 4c-d), which assumed no direct gene flow across the central Pyrenean axis and thus between the two most genetically differentiated lineages EPY and CWPY (Mantel’s R = 0.683, P = 0.001; Fig 6).

**Fig 6.**
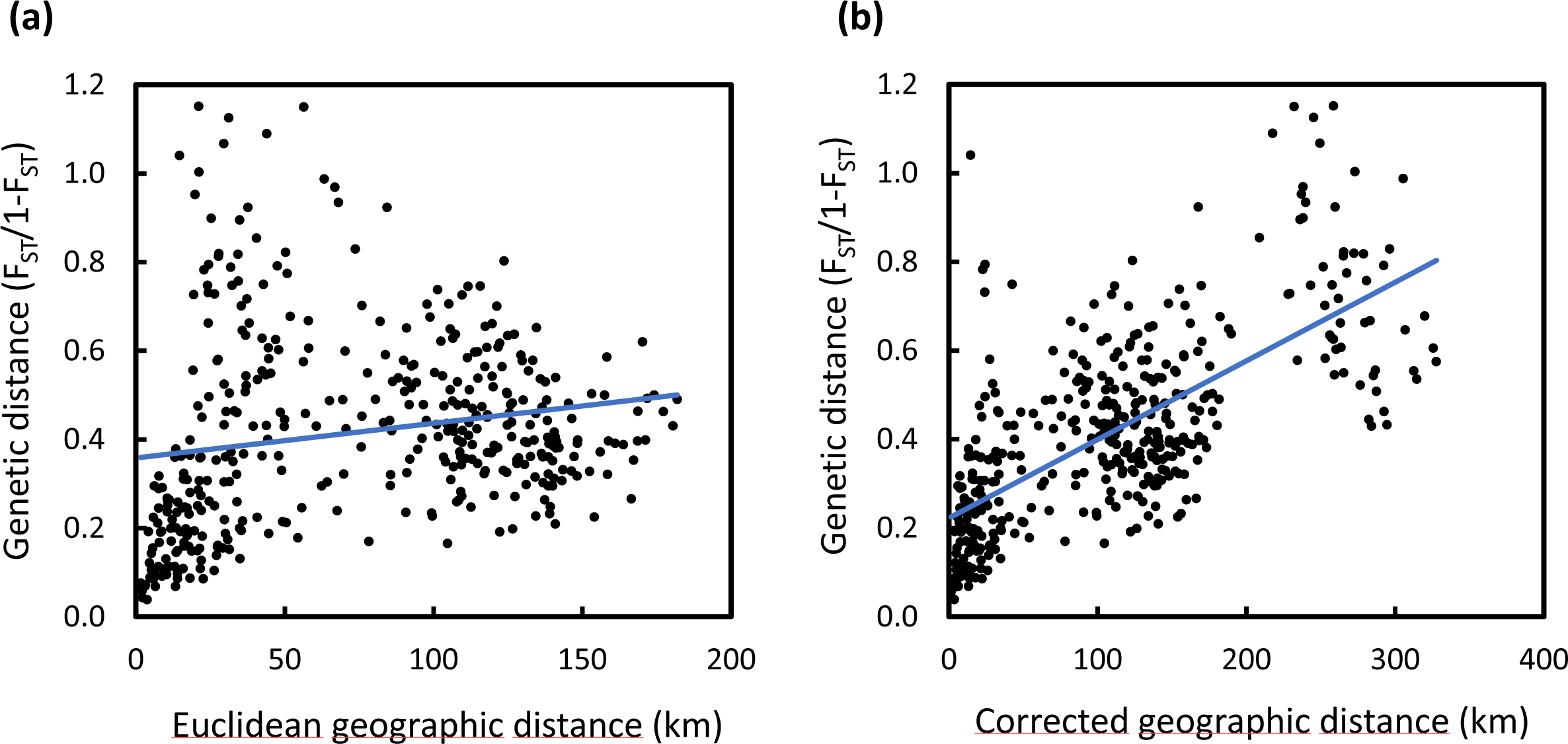
Isolation by distance analysis over *Alytes obstetricans*/*almogavarii* population pairs from the Pyrenees. Isolation by distance was calculated based on (a) Euclidean geographic distance between all pairs of populations (Mantel’s R = 0.185, P = 0.002) and (b) corrected geographic distance as suggested by PCA analysis, i.e. following a ring-shaped distribution around the Pyrenean chain (Fig 4; Mantel’s R = 0.683, P = 0.001).

The estimation of effective population sizes conducted in Colony returned, in general, low values (S1 Table). Values ranged from five in the population Laguna de Pájaros (LP) to 160 breeding individuals in the population Lago Ercina (ERC), with a mean Ne of 43. The lowest values were found in populations from the Guadarrama Mountains (Ne range = 5–15), while higher values were found in the Pyrenees (eastern Pyrenees: 14–51, central Pyrenees: 68, central-western Pyrenees: 12–84) and Picos de Europa (19–160).

### Demographic history

DIYABC analyses suggested highest support for two different but compatible scenarios of population divergence depending on the genetic markers used (i.e. scenario 2 when using only microsatellites and scenario 5 when combining microsatellites and ND4; Fig 2b, S5 Table). Both scenarios had non-overlapping 95% confidence intervals and type I and II errors for the best supported models were low (S5 Table), indicating high confidence in scenario choice. Model checking confirmed that the best supported scenarios provided a good fit to the observed data (data not shown). Furthermore, the mean mutation rate estimated for ND4 was found to be an order of magnitude higher than the literature value (1.96 x 10^-7^ substitutions/site/year; S6 Table), but still much lower than that estimated for microsatellites (ranging from 1.28 x 10^-4^ to 3.89 x 10^-4^ and from 4.59 x 10^-5^ to 3.31 x 10^-4^ for analyses conducted using only microsatellites and including both ND4 and microsatellite markers, respectively).

The analysis focused on microsatellites indicated that scenario 2 (the sequential splitting model based on results from clustering analyses) was the best supported model (Fig 2b, S5 Table). According to this scenario, the first split led to the separation of Pyrenean populations from peninsular ones. During the second split there was the divergence of, on the one hand, central-western and central-eastern Pyrenean populations, and on the other Picos de Europa and Guadarrama populations. Finally, populations from the central Pyrenees originated from eastern Pyrenean populations during the third split. The first split occurred 56 000–28 000 years ago, and the second and third splits 36 800–18 400 and 25 200–12 600 years ago, respectively (S6 Table). The analysis based on the combination of microsatellites and ND4, however, suggested that Picos de Europa populations were generated by admixture of populations from the central-western Pyrenees and Guadarrama (scenario 5; Fig 2b, S5 Table). Results indicated that the first split occurred 76 200–38 100 years ago, the second split 52 800–26 400 years ago and the admixture event 41 400–20 700 years ago (S6 Table).

## Discussion

The Iberian Peninsula is an extraordinary model system to assess the multiple historical vicariance events of species, the so called “refugia within refugia” model, where mountain ranges have had a significant role [11, 12]. Such processes are influenced by climatic and topographic conditions as well as through biological processes between populations, which in turn reflect different and contrasting time scales, historical and contemporary. To unravel such complex scenarios, a multilocus approach is expected to reveal contrasting, often discordant findings that underline intricate evolutionary processes [57]. Several recent studies targeting the Iberian Peninsula and the Mediterranean biodiversity hotspot have focused on the processes of niche divergence and admixture following secondary contact (e.g. Chamorro et al. [58] and Martínez-Freiría et al. [59] with Mediterranean vipers, Antunes et al. [60] with the fire salamander *Salamandra salamandra*, and Dufresnes et al. [61] with *Discoglossus* and *Pelodites* frogs), highlighting the role of these areas as centres of diversification and speciation. The results of the present study revealed a strong association of the defined genetic lineages with geography by using a multilocus analysis of genetic divergence in Iberian high mountain populations of *A. obstetricans*/*almogavarii*. Furthermore, high mountain populations showed higher levels of genetic distinctiveness than lowland populations (Fig 5), suggesting that mountains may have driven population differentiation through long-term geographic isolation, with lowlands more likely to be dispersal corridors for *A. obstetricans*/*almogavarii,* as it has been described for other species (see e.g. [62, 63]).

### The role of high mountains in genetic differentiation and glacial refugia

This study describes the presence of seven different nuclear microsatellite clusters, that are likely a result of recent microrefugial areas within the Pyrenees (Fig 5). Most of the recovered genetic structure was within Pyrenean populations, which in turn were genetically more related to each other than with the other two high mountain areas. Putative episodes of admixture within *A. obstetricans*/*almogavarii* in the Pyrenees were suggested by previous phylogenetic analyses [3]. Following the splits between major population lineages in *A. obstetricans*/*almogavarii*, which date back to the Early Pleistocene (starting from 2.5 Mya; [20]), there have been several glacial-interglacial episodes that likely provided opportunities for diverging taxa to come into contact and interbreed following range shifts tracking the climatic fluctuations [12, 64]. Our findings show that Pyrenean high mountain populations have gone through relatively recent events of admixture, likely favoured by the different glacial-interglacial episodes that characterised the Late Pleistocene.

Specifically, according to ABC analyses, the two lineages ascribed to *A. almogavarii* (E-F, central and eastern Pyrenees), together with some populations of *A. o. obstetricans* (central-western Pyrenean group) seem to have undergone a certain degree of contact and admixture until divergence took place ∼36 800–12 600 years ago (Fig 2b). Despite the selected model, which proposes two independent admixture events between the three lineages, the highly overlapping dates together with the significant ring-shaped isolation by distance pattern (Fig 6; see below) suggest that one single admixture event might have occurred within one main glacial refugia where the three lineages coincided. In addition, contemporary signs of connectivity between the two species were detected at lineage borders, as indicated by the occurrence of a contact zone between ND4 lineage E and *A. o. obstetricans* in the west (Figs 3, 5 and S2). Similarly, signs of admixture and extensive gene flow between *A. o. obstetricans* and *A. almogavarii* in the western Pyrenees were previously pointed out by allozyme markers [25, 26]. In contrast, in the central and eastern Pyrenees no signs of connectivity were detected between ND4 lineage F and either lineage E or *A. o. obstetricans*. This scenario of different contact zones telling different stories is exemplified in ring species, i.e. a system formed by a region of interconnected populations with both ends of the ring coming into contact without apparent admixture [65, 66]. In the Pyrenees, the spatial distribution of the *A. obstetricans* complex seems to fit with a postglacial colonisation pattern similar to what has been described in ring diversification, where taxa at the western part of the ring likely interbreed and those at the eastern side apparently don’t, displaying a continuum from slightly divergent neighbouring populations to substantially reproductively isolated taxa (Figs 4d, 5 and 6). Ring diversifications represent cases of population divergence around a geographical barrier in a ring-like manner [67] that can result in cases of speciation in action (i.e. ring species) and have been cited as evidence of evolution [68], with some of the most well-known cases being identified in herptiles, such as the *Ensatina eschscholtzii* salamander complex [69] and the western fence lizard *Sceloporus occidentalis* [70] in western North America. In light of this, our results are not entirely in line with the distinction of *A. almogavarii* as a different species, yet support a scenario of speciation in action, with lineage F showing evidence for speciation and reproductive isolation [23], and lineage E retaining some degree of connectivity with *A. o. obstetricans*. Nevertheless, we cannot rule out that this pattern may be the result of geographic sampling gaps, and additional surveys across hybrid zones between lineages E-F and *A. o. obstetricans*, as well as further work using resistance distances (i.e. by taking elevation, slope and land cover into account), would be required to fully test the ring species hypothesis.

Finally, with regard to the central-western Pyrenean group, the subdivision by microsatellite analyses into two or three different genetic clusters (Figs 4 and S2), points to a scenario of allopatric divergence after the recolonization of the Pyrenees as a result of geographic barriers, with consequent reduction or disruption of gene flow. Alternatively, this differentiation might have happened during one of the abrupt cooling episodes of the Holocene (e.g. the 8 200 year-event; [71]), with consequent isolation in separate glacial refugia. Range isolation and lineage divergence in separate Pyrenean refugia during Pleistocene glacial cycles were also invoked e.g. for the Pyrenean brook newt *Calotriton asper* [55], the ground-dwelling spider *Harpactocrates ravastellus* [72], the rusty-leaved alpenrose *Rhododendron ferrugineum* [73] and the snapdragon *Antirrhinum* [74], in line with evidence from a number of other high mountain areas that served as glacial refugia during periods of adverse conditions, such as the Andes, Himalaya and the Southern Alps in New Zealand (see [64] for a review). From a conservation point of view, we suggest that these areas should be treated as separate conservation and management units.

### Mito-nuclear discordances

The analyses of microsatellites distinguished two highly divergent *A. o. obstetricans* lineages, one in the central-western Pyrenees and the other in Picos de Europa (Figs 4 and S2), with the latter being more genetically related to the southern populations (ascribed to *A. o. pertinax*; Figs 4d and S7). In contrast, these two lineages bear the same mtDNA (ND4) haplogroup ascribed to *A. o. obstetricans* (haplogroup B; Figs 3 and S2a). One possible explanation is that the closer genetic affinity of the western and southern area populations rather than to the central- western Pyrenean lineage originated from extensive admixture with *A. o. obstetricans* (the central-western Pyrenean group) as the maternal donor and *A. o. pertinax* as the paternal donor (Fig 2b). A similar pattern was observed in *S. salamandra* [75] and in crustaceans of the genus *Daphnia* [76]. A plausible scenario could be that, during Late Pleistocene glacial periods (as suggested by results from DYABC modelling when combining both microsatellites and mtDNA; Fig 2), some *A. o. obstetricans* populations remained confined in isolated refugia where they coincided with *A. o. pertinax* for a sufficient time so that genetic admixture could take place. Similarly, range isolation during Pleistocene climatic fluctuations was suggested for the cryptic *Pelodytes* anuran clade [77, 78], the Cabrera vole *Microtus cabrerae* [79] and the scrub-legume grasshopper *Chorthippus binotatus binotatus* [80], where extant lineages likely diverged in separated refugia within the Iberian Peninsula. Subdivided glacial refugia could have experienced events of admixture during the succession of glacial and interglacial periods, with consequent fusion between refugial lineages [81, 82].

Mito-nuclear discordances with evidence for admixture or hybridisation are not uncommon in amphibians (e.g. [4, 8, 60]) and have also been described in a wide range of other animal species from north-central Iberian Peninsula, from arachnids [83] to mammals [84]. Furthermore, the Cantabrian Mountains represent a peculiar biogeographic region and a recognised hotspot of genetic diversity in amphibians, being home to endemic refugial clades in a number of species with broad European distribution [82, 85]. Finally, more recently, the Picos de Europa and *A. o. pertinax* populations may have come into secondary contact and interbred following an expansion phase, creating zones of admixture at lineage borders. Alternatively, the discordance may stem from stochastic processes in the form of genetic drift, and the phylogeographic structure detected in mtDNA (Figs 3 and S3) may have developed as a result of low individual dispersal distances and/or population sizes [86], as evidenced by our results and published literature (e.g. [87]). Further studies and sequencing of more molecular markers are needed to test these hypotheses and enrich our understanding of the phylogeography of *A. obstetricans*.

### Chytridiomycosis and population bottleneck

The genetic distinctiveness and limited genetic diversity of populations from the Guadarrama Mountains herein detected (Figs 4 and S2, Table 1) are likely early signs of inbreeding depression induced by an emerging pathogen, i.e. the chytrid fungus *Batrachochytrium dendrobatidis* [88]. It should be noted that our sampling was performed approximately 10 years after an epidemic of the disease chytridiomycosis, which hugely impacted this area in the period 1997–1999, causing several populations to decrease or even disappear [16, 89]. Accordingly, the only sampled population in the affected area that did not show signs of chytridiomycosis was the one presenting the highest genetic diversity (Montes de Valsain, MV; S1 Table). Our findings complement those of Albert et al. [18] that detected evidence of low genetic variability and strong population bottleneck in *A. obstetricans* from the same mountain system. In addition to this, we report on the first estimates of effective population sizes of these populations, which were among the lowest overall (Ne = 5–15; Table 1), raising concern for the long-term persistence of these populations, which are small and isolated. Indeed, the closest neighbouring populations are located more than 50 km away and are known to be small and not genetically related [18, 90]. Furthermore, the Guadarrama Mountains constitute a major barrier to gene flow in amphibians and a key feature shaping population structure and promoting population divergence across taxa [62]. Our results are an example of a disease acting as selective pressure in wild populations by inducing genetic hallmarks of bottlenecks and inbreeding, as it has also been shown in other species such as the black-tailed prairie dog (*Cynomys ludovicianus*), the mountain yellow-legged frog (*Rana muscosa*) and the bobcat (*Lynx rufus*) (reviewed in [91]).

### Differences in the evolutionary horizon/resolution between genetic markers

This study confirms previous findings in *A. obstetricans*/*almogavarii* clades [20] with the only difference that our species tree analyses recovered a different basal lineage (S3 Fig), a likely consequence of our limited inference power given that only two loci were used for species tree reconstruction. Results from ABC modelling based on either microsatellites or microsatellites + mtDNA resulted in time estimates highly different from those of the species tree (Fig 2b), suggesting that the two analyses are depicting different evolutionary events. In fact, while the species tree has proven useful to estimate divergence times associated with the formation of species/subspecies, ABC modelling was here employed to gain insights into the colonisation of three high mountain regions in the Iberian Peninsula by the different *A. obstetricans*/*almogavarii* taxa. Microsatellite markers, with their faster mutation rate, are known to perform poorly for the estimation of ancient historical events [57, 92]. On the contrary, microsatellites provide substantially better estimations than mtDNA for the most recent dynamics [57]. However, the combination of both markers in our ABC modelling resulted in similar divergence times as those estimated using microsatellites alone. It is thus possible that combining both markers may have biased toward microsatellites and the estimation of recent historical events.

We need to stress that DIYABC modelling has some uncertainty. Firstly, this approach is based on scenarios where no gene flow is permitted between populations after they initially diverge. Only single events of admixture between populations are considered, whereas recurrent gene flow due to dispersal cannot be incorporated. However, population structure analyses were used to investigate contemporary gene flow and identify patterns of admixture and contact zones at lineage borders (e.g. Fig 5). Secondly, the tested models do not represent a comprehensive range of all possible scenarios, but are instead based on a selection of contrasting hypotheses that we considered were most likely to reflect our data. We focused our analysis on simple contrasting models aimed at capturing the key demographic events, avoiding overcomplex and redundant models, therefore the selected models should be viewed as a starting point for evolutionary understanding. This approach has proven useful to increase the ability of DIYABC to reveal the correct model, as well as to better estimate the error and accuracy of parameter estimates [93]. On the other hand, it has to be noted that estimates based on mitochondrial data might be unreliable due to the erratic mutation rate of mtDNA and to the fact that mtDNA does not evolve neutrally since transmission of mitochondria is completely linked to maternal inheritance. Indeed, mtDNA-based estimates have been found to mismatch the onset of species divergence in both directions due to the stochasticity of coalescence [94].

### Concluding remarks

Our multilocus phylogeography across the Iberian Peninsula revealed high genetic structure correlated with geography and a complex pattern of lineage admixture in high mountain populations of *A. obstetricans*/*almogavarii*. Our study evidenced how each analysed mountain region underwent a peculiar phylogeographic history through the Late Pleistocene, which is consistent with the “refugia within refugia” model [11, 12] and confirms previous studies on a number of Iberian amphibian species (e.g. [22, 77, 85, 95, 96]). Results also support the assumption that refugia within refugia may be hotspots of extensive mito-nuclear discordances [82], highlighting the importance of multilocus approaches to infer the dynamics of population divergence. Environmental change in the different mountain systems may have an influence on selection, resulting in an increased divergence among isolated populations, consequently leading to speciation [97]. The phenetic differences between the subspecies of *Alytes* (especially *A. o. pertinax*) may be a further indication of adaptation to some micro-environmental differences such as streams vs still waters (known to influence coloration in *Alytes*, [98]), temperature and solar radiation at different altitudes between the studied mountain systems.

## Acknowledgements

We thank Saioa Fernández-Beaskoetxea, Amparo Mora, Susana Marquínez, Fernando Rivada, René A. Priego, Abel Bermejo and Matthew C. Fisher for field assistance, and Eva Albert, Laura Méndez, Annie Machordom and Andrés Fernández-Loras for laboratory assistance.

## Supporting information

**S1 Fig. Summary of *A. obstetricans*/*almogavarii* samples incorporated into each main analysis.** (a) Geographic location of populations included in each analysis. For population codes see Fig 1. The inset map shows the distribution of the main lineages: orange - ND4 haplogroups E-F (*A. almogavarii*), yellow - ND4 haplogroup B (*A. o. obstetricans*), blue - ND4 haplogroup A (*A. o. pertinax*), red - ND4 haplogroup C (*A. o. boscai*), green - ND4 haplogroup D (*A. o. boscai*), black - unclear (adapted from Dufresnes and Martínez-Solano [23]). (b) Markers used in each analysis, with the corresponding number of samples and populations. Effective population sizes were calculated only for populations with ≥ 15 genotyped individuals. In the case of demographic history, only high mountain populations were included in the analysis.

**S2 Fig. Results of phylogenetic and clustering analyses for *Alytes obstetricans*/*almogavarii*.** (a) Geographic distribution of the four mtDNA (ND4) haplogroups recovered in the analysis (blue: mtDNA haplogroup A, yellow: mtDNA haplogroup B, grey: mtDNA haplogroup E, orange: mtDNA haplogroup F). In four populations (T2, 61XR, BP and PI) we detected the presence of more than one haplogroup. (b) Results from Discriminant Analysis of Principal Components (DAPC) and (c) STRUCTURE for K = 7 microsatellite groups (see Fig 4). (d) Geographic distribution of the five genetic groups identified by microsatellite-based neighbour-joining analysis (see S7 Fig). White circles indicate populations with no data available for either ND4 or microsatellites. For population codes see Fig 1. The inset map shows the distribution of the main lineages: orange - ND4 haplogroups E-F (*A. almogavarii*), yellow - ND4 haplogroup B (*A. o. obstetricans*), blue - ND4 haplogroup A (*A. o. pertinax*), red - ND4 haplogroup C (*A. o. boscai*), green - ND4 haplogroup D (*A. o. boscai*), black - unclear (adapted from Dufresnes and Martínez-Solano [23]).

**S3 Fig. *Alytes* species tree produced in *BEAST based on one nuclear gene (β- fibint7) and four mitochondrial fragments (ND4, cyt-*b*, 12S and 16S).** Labels on branch tips correspond to the distinct ND4 haplogroups identified (blue: mtDNA haplogroup A, pink: mtDNA haplogroup B (central-western Pyrenean populations), yellow: mtDNA haplogroup B (Picos de Europa populations), grey: mtDNA haplogroup E, orange: mtDNA haplogroup F). Posterior probabilities of lineage divergence are indicated on branch labels.

**S4 Fig. Resulting plots from Discriminant Analysis of Principal Components (DAPC) across all *Alytes obstetricans*/*almogavarii* populations.** (a) Summary plots for K = 2-7 genetic clusters. At K = 2, genetic clusters are represented as density curves. At K = 3-7, dots represent individuals and genetic clusters are shown as inertia ellipses. Legend labels indicate the different genetic clusters: eastern Pyrenees (EPY, orange), central Pyrenees (CPY, grey), central-western Pyrenees (CWPY, pink), Picos de Europa mountains (PEU, yellow), and Guadarrama Mountain Range (GUA, blue). (b) Distribution of BIC (Bayesian Information Criterion) values according to the number of clusters. The red arrow indicates the number of clusters chosen for DAPC analysis. The optimal number of clusters was assessed using the *find.clusters* function and determined as the K value above which BIC (Bayesian Information Criterion) values decreased substantially.

**S5 Fig. Results of Bayesian clustering analyses in STRUCTURE for microsatellites.** (a) Mean (± SD) log probability of the data [Ln Pr(XΙK)] over 10 runs, for each value of K. (b) ΔK values as a function of K, calculated according to Evanno et al. [99].

**S6 Fig. Results of Tess3R analyses for microsatellites.** (a) Map depicting the distribution of ancestry coefficients inferred through Tess3R for K = 7 clusters. Black dots indicate the populations analysed. Colour codes as for STRUCTURE results. More saturated colours indicate a greater proportion of ancestry to either cluster. (b) Distribution of cross-validation scores according to the number of clusters. The red arrow indicates the number of clusters chosen for Tess3R analysis.

**S7 Fig. Neighbour-joining tree based on DA distances for microsatellite markers.** Branch colours delineate the seven genetic clusters inferred by STRUCTURE (see Figs 4a and 5), while colour shades around population codes correspond to the distinct mtDNA (ND4) haplogroups (blue: mtDNA haplogroup A, yellow: mtDNA haplogroup B, grey: mtDNA haplogroup E, orange: mtDNA haplogroup F; see Fig 3). In four populations (T2, 61XR, BP and PI) we detected the presence of more than one haplogroup. For population codes see S1 Table.

**S1 Table. Geographic information and standard genetic statistics of *Alytes obstetricans*/*almogavarii* sampling localities.** Populations are grouped according to the genetic group of interest (EPY: eastern Pyrenees, CPY: central Pyrenees, CWPY: central-western Pyrenees, PEU: Picos de Europa mountains, GUA: Guadarrama Mountain Range). Lat. – latitude, Long. – longitude, Alt. – altitude in meters, N – sample size for microsatellites, Na – mean number of alleles, Ar – allelic richness standardized for sample size, HO – observed heterozygosity, HE – expected heterozygosity, FIS – inbreeding coefficient, Ne – effective population size, N ND4 – sample size for ND4, ND4 haps – occurrence and code (in parentheses) of mitochondrial ND4 haplogroups identified in each population (see Fig 3), N cyt-b – sample size for cyt-*b*, N 12S – sample size for 12S, N 16S – sample size for 16S, N β- fibint7 – sample size for β-fibint7.

**S2 Table. Parameters used in DIYABC analysis and respective priors for the best supported scenarios.** The best supported scenarios were scenario 2 when considering only microsatellites (simple sequence repeats – SSRs) and scenario 5 when including both mtDNA (ND4) and microsatellite markers. See Fig 2 for more information on tested scenarios. N – effective population size for each analysed deme (EPY – eastern Pyrenees; CPY – central Pyrenees; CWPY – central-western Pyrenees; PEU – Picos de Europa Mountains, GUA – Guadarrama Mountains), ra –admixture rate, t – time of events in generations (t1 – time to the most recent split; t2 – time to the intermediate split; t3 – time to the most ancient split). Microsatellite (SSRs) and mitochondrial (ND4) parameters: mean *µ* – mean mutation rate, individual locus *µ* – individual locus mutation rate, mean *P* – mean coefficient *P*, individual locus *P* – individual locus coefficient *P*, SNI – Single Nucleotide Insertion rate, mean *k* – mean coefficient *k*, individual locus *k* – individual locus coefficient *k*. Microsatellite loci were divided in two groups depending on the motif length (tri- and tetranucleotide loci). Conditions: sequence data were simulated under a Tamura Nei (TN93) mutation model.

**S3 Table. Microsatellite-based (below diagonal) and ND4-based (above diagonal) pairwise estimates of FST between the seven genetic groups identified by STRUCTURE in *Alytes obstetricans*/*almogavarii* (see Fig 5).** All P values < 0.01.

**S4 Table. Analysis of molecular variance (AMOVA) for mitochondrial (ND4) and nuclear (microsatellites) markers based on the seven genetic groups identified by STRUCTURE in *Alytes obstetricans*/*almogavarii* (see Fig 5).** All P values < 0.001.

**S5 Table. Posterior probability of tested scenarios and 95% confidence intervals (CI) estimated with DIYABC analysis when considering only microsatellites and when including both mtDNA (ND4) and microsatellite markers.** Type I and II errors for the best supported scenarios are indicated. See Fig 2 for more information on tested scenarios.

**S6 Table. Posterior parameters (median and 95% confidence intervals) and RMedAD (Relative Median Absolute Deviation) estimated with DIYABC analysis for the best supported scenarios when considering only microsatellites (simple sequence repeats – SSRs; scenario 2) and when including both mtDNA (ND4) and microsatellite markers (scenario 5).** See Fig 2 for more information on tested scenarios. N – effective population size for each analysed deme (EPY – eastern Pyrenees; CPY – central Pyrenees; CWPY – central-western Pyrenees; PEU – Picos de Europa Mountains; GUA – Guadarrama Mountains), ra – admixture rate, t – time of events in generations (t1 – time to the most recent split; t2 – time to the intermediate split; t3 – time to the most ancient split), mean *µ* – mean mutation rate, mean *P* – mean coefficient *P*, mean *k* – mean coefficient *k, Q2.5* – quantile 2.5%, *Q97.5* – quantile 97.5%.

## S1 Appendix. Supporting methods.

Click here to access/download Supporting Information - Compressed/ZIP File Archive [Supporting information.zip](https://www.editorialmanager.com/pone/download.aspx?id=30607144&guid=a9522206-e0fc-4038-a243-ce2507fc9a6e&scheme=1)

