## Supplementary material for "Contrasting patterns of genetic admixture explain the phylogeographic history of Iberian high mountain populations of midwife toads": S1 Appendix.docx

**S1 Appendix. Supporting methods.**

*Nuclear and mitochondrial genes sequencing*

Primers used for amplification and sequencing were: for cyt-*b*, a shortened version of primers Ptacek1-L (5’-TGAGGACAAATATCATTCTGAGG-3’) [1] and CB3Xen-H (5’-GGCGAATAGGAARTATCATTC-3’) [2]; for 12S, primers 12Sa [3] and H1557mod [4]; for 16S, primers 16sar and 16Sbr [5]. Amplification conditions followed standard PCR amplifications with annealing temperatures ranging between 48-56 ºC. As for ND4, primers ND4 and Leu, described by Arevalo et al. [6], were initially used for amplification and sequencing. PCR conditions followed Gonçalves et al. [7]. However, we obtained non-resolvable electropherograms and/or very short sequences for a high number of individuals. We thus designed a new set of primers specific for *A. obstetricans*/*almogavarii*, ND4-F (5’-TACCCCTTTATTGGTCTCGC-3’) and Leu-F (5’-GGTTCCTAAGACCAACGGAT-3’), which allowed us to recover high quality sequences. PCR conditions were the same as described in Gonçalves et al. [7], except for the annealing temperature that was set to 57 ºC. Similarly, we designed specific primers for β-fibint7 based on sequences that amplified using β-fibint7 primers BFxF and BFxR [8]. These primers were AlyFibF (5’-GACCTTATGATAATCATCTTTATTGGCT-3’) and AlyFibR (5’-TGTTTGTTTGGATCTGCATGTAGCCTGG-3’), with annealing temperatures ranging between 50-56 ºC. Primer combinations were: BFxF-BFxR, BFxF-AlyFibR and AlyFibF-BFxR.

*Genetic structure*

For microsatellites, population structure was inferred with a Discriminant Analysis of Principal Components (DAPC), using the ADEGENET package 2.1.1 [9, 10] in R 3.5.1 [11] (S1 Fig). The optimal number of clusters was assessed using the *find.clusters* function and determined as the K value above which BIC (Bayesian Information Criterion) values decreased substantially. We covered a range of possible clusters from 1 to 100 and retained all principal components (PCs), as suggested by Jombart and Collins [12]. We also used DAPC to investigate the hierarchical genetic structure of our data through the examination of various values of K and the sequence of differentiation when K increases. The same package was also used to perform a Principal Component Analysis (PCA), for comparison purposes. For DAPC analysis, 150 PCs (80% of variance) and all discriminant functions were retained. The number of PCs retained for DAPC was estimated using cross-validation through the *xvalDapc* command. Furthermore, the Bayesian cluster analysis implemented in STRUCTURE 2.3.4 [13] was performed to corroborate the optimal clustering solution inferred by DAPC and PCA analyses. All runs were repeated 10 times for each K, set between 1 and 15, with 100K burn-in steps followed by 100K MCMC repetitions. We used the admixture model with correlated allele frequencies. The optimal number of genetic clusters was determined using both the original method of Pritchard et al. [13] and the ΔK method of Evanno et al. [14], as implemented in STRUCTURE HARVESTER 0.6.94 [15]. The R package pophelper [16] was used to average replicate runs of the optimal K [17] and plot the final output. Genetic relationships between STRUCTURE clusters were visualised by constructing a neighbour-joining (NJ) tree based on net nucleotide distances [18] using the program NEIGHBOR in the PHYLIP package 3.695 [19]. In addition, to visualise genetic divergence between sampled sites and check for consistency with ND4-inferred lineages, we drew a NJ tree using the program POPTREEW [20]. We used Nei’s genetic distance (D_A_, [21]) and performed 1 000 bootstraps. As POPTREEW does not allow loci to have no data for an entire locality, we excluded 22 localities with data missing for at least one locus. Finally, we complemented previous analyses with a spatial-based clustering approach implemented in the R package Tess3R 1.1.0, a spatially explicit least-squares optimisation approach that incorporates geographic proximity information and a model-free algorithm [22]. Tess3R was run with 10 replicates for each of K=1-50 using default parameters, and the optimal K value was chosen using the cross-validation score as the value of K that corresponded to a plateau of the curve.

*Demographic history (DIYABC)*

To reduce computational demands, we selected 50 individuals from each of the five population groups, maximizing the number of samples with available ND4 sequences and minimizing missing data for microsatellites. The mutation rate prior distribution assumed for ND4 included a range of values comprehensive of the mutation rate calculated in a previous phylogenetic study (0.85 x 10^-8^ substitutions/site/year; [23]). We also performed a preliminary analysis run using a fixed prior for mutation rate that appeared unsuccessful, as the estimation of prior distribution of parameters showed a lack of correspondence between simulated and observed data sets (data not shown). The final parameter setting is shown in S2 Table. We generated 10^6^ simulated datasets per scenario, assuming a 1:1 female to male sex ratio and a generation time of 1 to 2 years [24]. The following summary statistics were used for microsatellites: mean number of alleles, mean genetic diversity and mean allele size variance as one sample statistics, and Fst and (dµ)² distance as pairwise statistics. As for ND4, the following summary statistics were used: number of segregating sites, mean of pairwise differences, variance of pairwise differences, Tajima’s D and private segregating sites as one sample statistics, and number of haplotypes and Fst as pairwise statistics. Pre-evaluation of scenarios, selection of the most supported scenario, confidence in scenario choice (type I and II errors), model checking, estimation of the posterior distribution of parameters and evaluation of bias and precision on parameters estimation for the most supported scenario followed Lucati et al. [25].
