## Supplementary material for "Contrasting patterns of genetic admixture explain the phylogeographic history of Iberian high mountain populations of midwife toads": S1 Fig.pdf

(a)

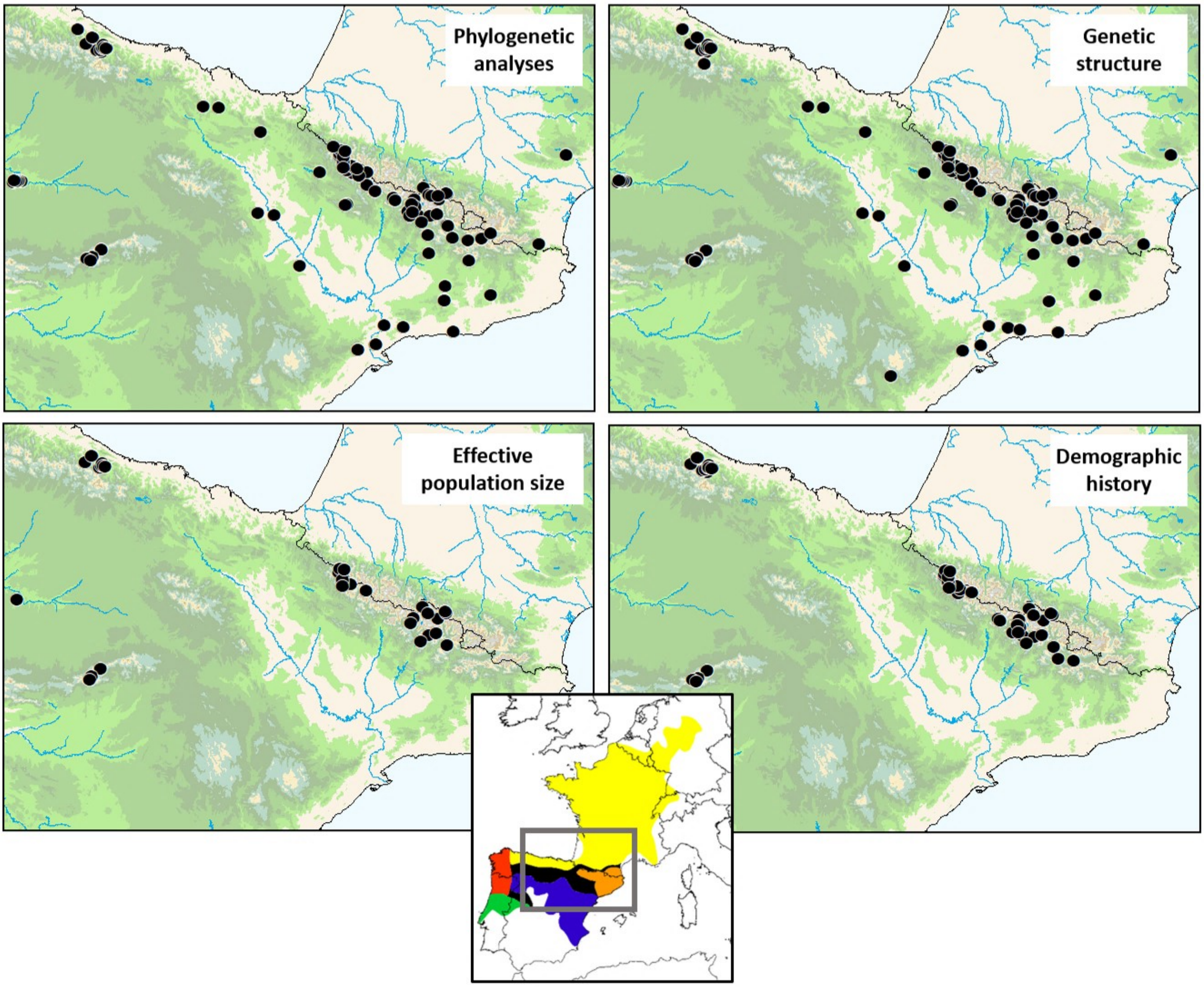


(b)

| Type of analysis | Markers used | N Samples/Populations |
|---|---|---|
| Phylogenetic analyses | Nuclear (β-fibint7) and mitochondrial (ND4, cyt-*b*, 12S, 16S) gene fragments | β-fibint7: 20/15, ND4: 219/95, cyt-*b*: 40/26, 12S: 42/28, 16S: 40/27 |
| Genetic structure | Microsatellites | 878/102 |
| Effective population size | Microsatellites | 622/34 |
| Demographic history | Microsatellites + ND4 | Microsatellites: 250/55, ND4: 136/55 |
