## Supplementary material for "Contrasting patterns of genetic admixture explain the phylogeographic history of Iberian high mountain populations of midwife toads": S1 Table.docx

**S1 Table. Geographic information and standard genetic statistics of *Alytes obstetricans*/*almogavarii* sampling localities.** Populations are grouped according to the genetic group of interest (EPY: eastern Pyrenees, CPY: central Pyrenees, CWPY: central-western Pyrenees, PEU: Picos de Europa mountains, GUA: Guadarrama Mountain Range).

| Genetic group | Population | Code | Lat. | Long. | Alt. | N | Na | Ar | Ho | He | Fis | N_e_ | N ND4 | ND4 haps | N  cyt-*b* | N 12S | N 16S | N  β-fibint7 |
| --- | --- | --- | --- | --- | --- | --- | --- | --- | --- | --- | --- | --- | --- | --- | --- | --- | --- | --- |
| EPY_mountains_ | Abeurador Abella | Ab | 42.452 | 0.564 | 1360 | 3 | - | - | - | - | - | - | 1 | 1(F) | - | - | - | - |
|  | Abeurador Boscalt | AB | 42.314 | 1.587 | 1450 | 1 | - | - | - | - | - | - | 2 | 2(F) | - | - | - | - |
|  | Abeurador Sarroqueta | AS | 42.444 | 0.724 | 1106 | 3 | - | - | - | - | - | - | 2 | 2(F) | - | - | - | - |
|  | Abeurador Señuy | Sen | 42.460 | 0.641 | 1227 | 5 | 2.412 | 1.440 | 0.388 | 0.392 | 0.126 | - | 2 | 2(F) | - | - | - | - |
|  | Barranc de Viu | BV | 42.373 | 0.812 | 1125 | 19 | 6.353 | 1.630 | 0.471 | 0.606 | 0.279 | 14 | 2 | 2(F) | - | - | - | - |
|  | Barranco Llana de Obarra | BLO | 42.528 | 0.653 | 1421 | 5 | 3.647 | 1.620 | 0.535 | 0.547 | 0.158 | - | 2 | 2(F) | - | - | - | - |
|  | Bassa Manyanet | BM | 42.470 | 0.908 | 2277 | 19 | 4.765 | 1.590 | 0.53 | 0.571 | 0.104 | 33 | 1 | 1(F) | - | - | - | - |
|  | Bassa Vallibierna | Bas | 42.613 | 0.610 | 1912 | 19 | 3.294 | 1.480 | 0.432 | 0.468 | 0.104 | 26 | 3 | 3(F) | - | - | - | - |
|  | Bellver de Cerdanya | 15 | 42.372 | 1.777 | 1087 | 1 | - | - | - | - | - | - | 1 | 1(F) | - | - | - | - |
|  | Enveitg | S50 | 42.459 | 1.906 | 1537 | 3 | - | - | - | - | - | - | 3 | 3(F) | - | - | - | - |
|  | Estany Basibé | EB | 42.548 | 0.596 | 2254 | 19 | 5.412 | 2.690 | 0.494 | 0.595 | 0.198 | 53 | 2 | 2(F) | - | - | - | - |
|  | Estanyet Coma d'Espós | ECE | 42.509 | 1.017 | 2406 | 18 | 5.235 | 1.60 | 0.508 | 0.573 | 0.149 | 51 | 1 | 1(F) | - | - | - | - |
|  | Estanyet de Davall | ED | 42.415 | 1.232 | 2058 | 18 | 6.059 | 2.740 | 0.494 | 0.647 | 0.265 | 51 | 3 | 3(F) | - | - | - | - |
|  | Les Paüls | S88 | 42.408 | 0.606 | 1242 | 2 | - | - | - | - | - | - | 1 | 1(F) | - | - | - | - |
|  | Navès | S3 | 42.088 | 1.677 | 1010 | 1 | - | - | - | - | - | - | 1 | 1(F) | - | - | - | - |
|  | Safareig de Taüll | ST | 42.522 | 0.846 | 1515 | 7 | 2.706 | 2.550 | 0.366 | 0.425 | 0.230 | - | - | - | - | - | - | - |
|  | Serra de l’Orri | 65 | 42.428 | 1.205 | 1592 | 1 | - | - | - | - | - | - | 1 | 1(F) | 2 | 2 | 2 | 1 |
| EPY_lowland_ | Castellterçol | Aly13 | 41.763 | 2.118 | 678 | 1 | - | - | - | - | - | - | 1 | 1(F) | 1 | 1 | 1 | - |
|  | Els Banys i Palaldà | M14 | 42.473 | 2.670 | 305 | 1 | - | - | - | - | - | - | - | - | 1 | 1 | 1 | 1 |
|  | Ferran | Aly12 | 41.730 | 1.401 | 672 | - | - | - | - | - | - | - | 1 | 1(F) | 1 | 1 | 1 | - |
|  | Gavet de la Conca | 77 | 42.045 | 1.037 | 948 | 2 | - | - | - | - | - | - | 2 | 2(F) | 1 | 1 | 1 | - |
|  | La Goda | Aly8 | 41.563 | 1.446 | 691 | 1 | - | - | - | - | - | - | 2 | 2(F) | 4 | 4 | 4 | - |
|  | La Pobla de Segur | S16 | 42.250 | 0.963 | 812 | 1 | - | - | - | - | - | - | 1 | 1(F) | - | - | - | - |
|  | Noves de Segre | NS | 42.295 | 1.342 | 713 | 11 | 4.294 | 2.680 | 0.447 | 0.564 | 0.258 | - | 2 | 2(F) | - | - | - | - |
|  | Pobla de Carivenys | Aly3 | 41.570 | 1.441 | 691 | 3 | - | - | - | - | - | - | 2 | 2(F) | 1 | 1 | 1 | 1 |
|  | Vilanova i la Geltrú | M26 | 41.239 | 1.683 | 62 | 1 | - | - | - | - | - | - | - | - | 3 | 3 | 3 | 1 |
| CPY_mountains_ | Aljibe Pino Simón | APS | 42.561 | 0.291 | 1426 | 4 | - | - | - | - | - | - | 2 | 2(E) | - | - | - | - |
|  | Arguis | N67 | 42.326 | 0.413 | 1211 | 2 | - | - | - | - | - | - | 4 | 4(E) | 2 | 2 | 2 | 2 |
|  | Balsa Pertacua | BP | 42.714 | 0.423 | 1913 | 7 | 3.588 | 1.540 | 0.479 | 0.504 | 0.132 | - | 2 | 1(B), 1(E) | - | - | - | - |
|  | Bassa de la Mora | Mor | 42.545 | 0.326 | 1903 | 14 | 4.059 | 5.350 | 0.272 | 0.407 | 0.403 | - | 3 | 3(E) | - | - | - | - |
|  | Buisán | XR18 | 42.585 | 0.010 | 1482 | 1 | - | - | - | - | - | - | 1 | 1(E) | - | - | - | - |
|  | Ibón de los Asnos | 61XR | 42.693 | 0.267 | 2003 | 4 | - | - | - | - | - | - | 4 | 2(B), 2(E) | - | - | - | - |
|  | Ibón Serrato Bajo | ISB | 42.765 | 0.215 | 2447 | 3 | - | - | - | - | - | - | 2 | 2(B) | - | - | - | - |
|  | Linás de Broto | XR16 | 42.623 | 0.168 | 1504 | 1 | - | - | - | - | - | - | 1 | 1(E) | - | - | - | - |
|  | Monrepos | M3Q | 42.347 | 0.391 | 1214 | 1 | - | - | - | - | - | - | - | - | 1 | 1 | 1 | 1 |
|  | Plano de Igüer | PI | 42.745 | 0.588 | 1585 | 18 | 6.588 | 1.650 | 0.447 | 0.629 | 0.318 | 68 | 2 | 1(B), 1(E) | - | - | - | - |
| CPY_lowland_ | Asó-Veral | N63 | 42.610 | 0.925 | 567 | 2 | - | - | - | - | - | - | 2 | 2(E) | 1 | 1 | 1 | 1 |
| CWPY_mountains_ | Arties | M7 | 42.699 | 0.862 | 1548 | 3 | - | - | - | - | - | - | - | - | 1 | 2 | - | - |
|  | Barranco Las Foyas | BLF | 42.862 | 0.696 | 1372 | 4 | - | - | - | - | - | - | 2 | 2(B) | - | - | - | - |
|  | Bassa d'Arres | BA | 42.769 | 0.715 | 1562 | 19 | 2.765 | 5.620 | 0.338 | 0.336 | 0.023 | 21 | 4 | 4(B) | - | - | - | - |
|  | Canfranc | CAN | 42.717 | 0.523 | 1501 | 2 | - | - | - | - | - | - | 1 | 1(B) | 2 | 2 | 2 | 1 |
|  | Estanhet d'Arcoïls | Arc | 42.680 | 0.988 | 2392 | 20 | 2.235 | 1.270 | 0.271 | 0.262 | 0.036 | 20 | 3 | 3(B) | - | - | - | - |
|  | Estanho Vilac | EV | 42.709 | 0.814 | 1638 | 20 | 2.059 | 5.510 | 0.197 | 0.222 | 0.140 | 21 | 2 | 2(B) | - | - | - | - |
|  | Estany d'Aulà | EA | 42.769 | 1.099 | 2128 | 20 | 2.471 | 1.360 | 0.292 | 0.352 | 0.197 | 17 | 3 | 3(B) | - | - | - | - |
|  | Estany de Clavera | ECl | 42.778 | 1.077 | 2230 | 20 | 2.353 | 1.340 | 0.308 | 0.335 | 0.106 | 25 | 3 | 3(B) | - | - | - | - |
|  | Formigal | XR27 | 42.775 | 0.376 | 1823 | 3 | - | - | - | - | - | - | 3 | 3(B) | - | - | - | - |
|  | Ibón Acherito | IA | 42.880 | 0.707 | 1872 | 21 | 5.588 | 1.560 | 0.493 | 0.548 | 0.127 | 51 | 5 | 5(B) | 2 | 1 | 1 | 1 |
|  | Ibón Campo de Troya Inferior | ICT | 42.766 | 0.409 | 2071 | 10 | 4.294 | 1.530 | 0.400 | 0.499 | 0.249 | - | 3 | 3(B) | - | - | - | - |
|  | Ibón Espelunciecha | IBE | 42.787 | 0.431 | 1951 | - | - | - | - | - | - | - | 1 | 1(B) | - | - | - | - |
|  | Ibón Negras | IN | 42.787 | 0.465 | 2079 | 18 | 3.941 | 1.450 | 0.378 | 0.433 | 0.155 | 44 | 4 | 4(B) | - | - | - | - |
|  | Ibón Orná | IO | 42.798 | 0.614 | 1851 | 5 | 2.941 | 1.470 | 0.379 | 0.422 | 0.213 | - | 3 | 3(B) | - | - | - | - |
|  | Ibón Serrato Alto | ISA | 42.765 | 0.213 | 2459 | 17 | 3.529 | 1.460 | 0.332 | 0.443 | 0.286 | 27 | 4 | 4(B) | - | - | - | - |
|  | Ibón Viejo | IV | 42.781 | 0.602 | 2111 | 20 | 5.588 | 1.560 | 0.400 | 0.550 | 0.296 | 22 | 2 | 2(B) | - | - | - | - |
|  | Isaba | N85 | 42.946 | 0.835 | 1227 | 2 | - | - | - | - | - | - | 2 | 2(B) | 1 | 1 | 1 | 1 |
|  | Lac d'Arlet | A | 42.839 | 0.615 | 1998 | 15 | 5.176 | 1.550 | 0.379 | 0.525 | 0.316 | 84 | 2 | 2(B) | - | - | - | - |
|  | Lac de Lhurs | L | 42.922 | 0.704 | 1698 | 15 | 4.176 | 1.530 | 0.422 | 0.509 | 0.209 | 52 | 2 | 2(B) | - | - | - | - |
|  | Lescun | V | 42.934 | 0.637 | 903 | 15 | 3.353 | 1.480 | 0.345 | 0.459 | 0.286 | 12 | 3 | 3(B) | - | - | - | - |
|  | Naval Aguas Tuertas | NAT | 42.812 | 0.621 | 1623 | 19 | 6.118 | 1.630 | 0.501 | 0.613 | 0.213 | 49 | 3 | 3(B) | - | - | - | - |
|  | Pla de Beret | M9 | 42.725 | 0.964 | 2070 | 1 | - | - | - | - | - | - | 1 | 1(B) | 1 | 1 | 1 | - |
|  | Puit d'Arious | P | 42.864 | 0.633 | 1874 | 11 | 4.588 | 1.570 | 0.437 | 0.53 | 0.249 | - | 2 | 2(B) | - | - | - | - |
|  | Tramacastilla de Tena | XR23 | 42.705 | 0.320 | 1267 | 1 | - | - | - | - | - | - | 1 | 1(B) | - | - | - | - |
| CWPY_lowland_ | Lac du Saut de Vésoles | M15 | 43.555 | 2.794 | 992 | 1 | - | - | - | - | - | - | 1 | 1(B) | 1 | 1 | 1 | 1 |
| PEU_mountains_ | Ándara | AND | 43.213 | 4.716 | 1346 | 14 | 4.941 | 1.670 | 0.519 | 0.648 | 0.235 | - | 1 | 1(B) | - | - | - | - |
|  | Ftes. Carrionas | Leo | 43.011 | 4.744 | 2079 | 2 | - | - | - | - | - | - | - | - | - | - | - | - |
|  | Lago de Valdominguero | Val | 43.207 | 4.727 | 1834 | 19 | 5.647 | 1.650 | 0.592 | 0.635 | 0.097 | 26 | 3 | 3(B) | - | - | - | - |
|  | Lago Ercina | ERC | 43.267 | 4.979 | 1120 | 16 | 7.824 | 1.720 | 0.652 | 0.697 | 0.098 | 160 | 2 | 2(B) | - | - | - | - |
|  | Liordes | Li | 43.149 | 4.857 | 1885 | 14 | 4.588 | 1.620 | 0.562 | 0.598 | 0.100 | - | 3 | 3(B) | - | - | - | - |
|  | Llagos de Jesús | LJ | 43.175 | 5.056 | 1199 | 19 | 5.824 | 1.670 | 0.602 | 0.654 | 0.108 | 98 | 2 | 2(B) | - | - | - | - |
|  | Pilón de Igüedri | IGU | 43.146 | 4.774 | 1386 | 10 | 4.706 | 1.730 | 0.587 | 0.687 | 0.200 | - | 3 | 3(B) | - | - | - | - |
|  | Pilón de Moñetas | PMON | 43.202 | 4.783 | 1530 | 20 | 4.647 | 1.670 | 0.602 | 0.654 | 0.106 | 19 | 3 | 3(B) | - | - | - | - |
|  | Pilón de Pandébano | PP | 43.235 | 4.781 | 1249 | 18 | 6.471 | 1.740 | 0.602 | 0.720 | 0.195 | 87 | 3 | 3(B) | - | - | - | - |
|  | Pilón Vegas de Sotres | PVS | 43.208 | 4.767 | 1530 | 18 | 6.471 | 1.730 | 0.635 | 0.711 | 0.136 | 20 | 3 | 3(B) | - | - | - | - |
|  | Pilón Vegas del Enol | PVE | 43.269 | 4.998 | 1120 | 18 | 7.824 | 1.710 | 0.682 | 0.688 | 0.038 | 77 | 3 | 3(B) | - | - | - | - |
|  | Pozo de Moñetas | MON | 43.197 | 4.786 | 1530 | 17 | 4.353 | 1.440 | 0.419 | 0.430 | 0.058 | 32 | 3 | 3(B) | - | - | - | - |
|  | Pozos de Lloroza | LLO | 43.165 | 4.811 | 1482 | 17 | 5.824 | 1.710 | 0.653 | 0.692 | 0.087 | 68 | 2 | 2(B) | - | - | - | - |
| PEU_lowland_ | Murua | Mur | 42.976 | 2.736 | 638 | 1 | - | - | - | - | - | - | 1 | 1(B) | - | - | - | - |
|  | Puerto de Lizarraga | M10 | 42.860 | 2.005 | 928 | 1 | - | - | - | - | - | - | 1 | 1(B) | 1 | 1 | 1 | - |
|  | Valle del Tendi | M2 | 43.302 | 5.249 | 464 | 1 | - | - | - | - | - | - | 1 | 1(B) | 1 | 1 | 1 | 1 |
| GUA_mountains_ | Charcas de la Rubia | CHR | 40.846 | 3.949 | 1898 | 18 | 3.471 | 1.490 | 0.475 | 0.474 | 0.029 | 11 | 3 | 3(A) | - | - | - | - |
|  | Charcas del Salto | ES | 40.846 | 3.948 | 1898 | 11 | 3.176 | 1.480 | 0.463 | 0.451 | 0.030 | - | 3 | 3(A) | - | - | - | - |
|  | Charcas Secas | CHS | 40.848 | 3.948 | 1898 | 8 | 2.412 | 1.380 | 0.385 | 0.357 | 0.007 | - | 3 | 3(A) | - | - | - | - |
|  | Circo del Nevero | CN | 40.979 | 3.844 | 1980 | 20 | 2.941 | 1.440 | 0.377 | 0.425 | 0.152 | 15 | 3 | 3(A) | - | - | - | - |
|  | Laguna de Pájaros | LP | 40.860 | 3.948 | 1898 | 20 | 3.471 | 1.490 | 0.430 | 0.476 | 0.124 | 5 | 3 | 3(A) | - | - | - | - |
|  | Laguna Grande | LG | 40.840 | 3.957 | 1898 | 1 | - | - | - | - | - | - | 1 | 1(A) | - | - | - | - |
|  | Montes de Valsaín | MV | 40.824 | 4.017 | 1527 | 11 | 3.765 | 1.580 | 0.514 | 0.558 | 0.125 | - | 3 | 3(A) | - | - | - | - |
|  | Puerto de Cotos | C | 40.823 | 3.962 | 1829 | 5 | 3.000 | 1.560 | 0.485 | 0.501 | 0.144 | - | 3 | 3(A) | - | - | - | - |
|  | Valdesqui | VQ | 40.815 | 3.961 | 1829 | 18 | 4.588 | 1.530 | 0.431 | 0.510 | 0.188 | 20 | 3 | 3(A) | - | - | - | - |
| GUA_lowland_ | Barranc de la Galera | N50 | 40.746 | 0.332 | 723 | 1 | - | - | - | - | - | - | - | - | 1 | 1 | 1 | - |
|  | Bassa de Saranou | M21 | 40.864 | 0.581 | 243 | 1 | - | - | - | - | - | - | 1 | 1(A) | 1 | 1 | 1 | - |
|  | Bassa Gran | M20 | 40.860 | 0.571 | 243 | - | - | - | - | - | - | - | 1 | 1(A) | 1 | 1 | 1 | - |
|  | Buñuel | Bun | 41.982 | 1.438 | 240 | 3 | - | - | - | - | - | - | 5 | 5(A) | 3 | 3 | 3 | 3 |
|  | Fte. de la Marlota | T10 | 41.377 | 5.452 | 751 | 2 | - | - | - | - | - | - | 4 | 4(A) | - | - | - | - |
|  | Fte. de los Perros | T6 | 41.395 | 5.423 | 693 | 5 | 4.000 | 1.690 | 0.659 | 0.615 | 0.056 | - | 4 | 4(A) | - | - | - | - |
|  | Fte. de Picarico | T7 | 41.400 | 5.451 | 751 | 6 | 4.471 | 1.720 | 0.794 | 0.651 | 0.110 | - | 4 | 4(A) | - | - | - | - |
|  | Fte. Los Billares | T5 | 41.409 | 5.393 | 772 | 15 | 6.824 | 1.750 | 0.646 | 0.719 | 0.138 | 70 | 3 | 3(A) | - | - | - | - |
|  | Fte. Nueva de Bardales | T2 | 41.424 | 5.344 | 706 | 5 | 4.471 | 1.660 | 0.562 | 0.590 | 0.160 | - | 4 | 2(A), 2(B) | - | - | - | - |
|  | Fte. Valdespino | T4 | 41.416 | 5.387 | 751 | 4 | - | - | - | - | - | - | 4 | 4(A) | - | - | - | - |
|  | L'Argentera | Aly7 | 41.144 | 0.921 | 341 | 2 | - | - | - | - | - | - | - | - | - | 1 | 1 | - |
|  | Mas de Barberans | Aly5 | 40.748 | 0.325 | 723 | 1 | - | - | - | - | - | - | 2 | 2(A) | 2 | 2 | 2 | - |
|  | Mora la Nova | 1012 | 41.109 | 0.642 | 52 | 3 | - | - | - | - | - | - | 1 | 1(A) | - | 1 | 1 | - |
|  | Nacimiento Nervión | M12 | 42.939 | 2.982 | 830 | 1 | - | - | - | - | - | - | 1 | 1(A) | 2 | 2 | 2 | 2 |
|  | Nogueruelas | TER | 40.235 | 0.633 | 1203 | 1 | - | - | - | - | - | - | - | - | - | - | - | - |
|  | Novallas | NOV | 41.947 | 1.950 | 435 | 3 | - | - | - | - | - | - | 2 | 2(A) | - | - | - | - |
|  | Reus | M5 | 41.150 | 1.106 | 108 | 1 | - | - | - | - | - | - | - | - | - | - | - | - |
|  | Torrecilla de Valmadrid | Tor | 41.501 | 0.854 | 396 | 3 | - | - | - | - | - | - | 2 | 2(A) | 2 | 2 | 2 | 2 |
|  | Total |  |  |  |  | 878 |  |  |  |  |  |  | 219 |  | 40 | 42 | 40 | 20 |

Lat. – latitude, Long. – longitude, Alt. – altitude in meters, N – sample size for microsatellites, Na – mean number of alleles, Ar – allelic richness standardized for sample size, H_O_ – observed heterozygosity, H_E_ – expected heterozygosity, F_IS_ – inbreeding coefficient, N_e_ – effective population size, N ND4 – sample size for ND4, ND4 haps – occurrence and code (in parentheses) of mitochondrial ND4 haplogroups identified in each population (see Fig 3), N cyt-*b* – sample size for cyt-*b*, N 12S – sample size for 12S, N 16S – sample size for 16S, N β-fibint7 – sample size for β-fibint7.
