## Supplementary material for "Contrasting patterns of genetic admixture explain the phylogeographic history of Iberian high mountain populations of midwife toads": S2 Table.docx

**S2 Table. Parameters used in DIYABC analysis and respective priors for the best supported scenarios.** The best supported scenarios were scenario 2 when considering only microsatellites (simple sequence repeats – SSRs) and scenario 5 when including both mtDNA (ND4) and microsatellite markers. See Fig 2 for more information on tested scenarios.

|  | Microsatellites | | | | Microsatellites + ND4 | |
| --- | --- | --- | --- | --- | --- | --- |
| Parameter | | Conditions | Distribution [min-max] | Conditions | | Distribution [min-max] |
| N_EPY_ | |  | Uniform [10 - 20 000] |  | | Uniform [10 - 20 000] |
| N_CPY_ | |  | Uniform [10 - 20 000] |  | | Uniform [10 - 20 000] |
| N_CWPY_ | |  | Uniform [10 - 20 000] |  | | Uniform [10 - 20 000] |
| N_PEU_ | |  | Uniform [10 - 20 000] |  | | Uniform [10 - 20 000] |
| N_GUA_ | |  | Uniform [10 - 20 000] |  | | Uniform [10 - 20 000] |
| N_EPY-GUA_ | |  | Uniform [10 - 40 000] |  | | - |
| N_EPY-CWPY-GUA_ | |  | - |  | | Uniform [10 - 40 000] |
| ra | |  | - |  | | 0.001 – 0.999 |
| t_1_ | |  | Uniform [10 - 30 000] |  | | Uniform [10 - 30 000] |
| t_2_ | | t_2_>t_1_ | Uniform [10 - 40 000] | t_2_>t_1_ | | Uniform [10 - 40 000] |
| t_3_ | | t_3_>t_2_ | Uniform [10 - 50 000] | t_3_>t_2_ | | Uniform [10 - 50 000] |
| Mean *µ1*_(SSRs)_ | |  | Uniform [10^-5^ - 10^-3^] |  | | Uniform [10^-5^ - 10^-3^] |
| Individual locus *µ1*_(SSRs)_ | |  | Gamma [10^-5^ - 10^-2^] |  | | Gamma [10^-5^ - 10^-2^] |
| Mean *P1*_(SSRs)_ | |  | Uniform [10^-1^ - 3x10^-1^] |  | | Uniform [10^-1^ - 3x10^-1^] |
| Individual locus *P1*_(SSRs)_ | |  | Gamma [10^-2^ - 9x10^-1^] |  | | Gamma [10^-2^ - 9x10^-1^] |
| SNI_(SSRs)_ | |  | Log-u [0] |  | | Log-u [0] |
| Mean *µ2*_(SSRs)_ | |  | Uniform [10^-5^ - 10^-3^] |  | | Uniform [10^-5^ - 10^-3^] |
| Individual locus *µ2*_(SSRs)_ | |  | Gamma [10^-5^ - 10^-2^] |  | | Gamma [10^-5^ - 10^-2^] |
| Mean *P2*_(SSRs)_ | |  | Uniform [10^-1^ - 3x10^-1^] |  | | Uniform [10^-1^ - 3x10^-1^] |
| Individual locus *P2*_(SSRs)_ | |  | Gamma [10^-2^ - 9x10^-1^] |  | | Gamma [10^-2^ - 9x10^-1^] |
| SNI_(SSRs)_ | |  | Log-u [0] |  | | Log-u [0] |
| Mean *µ*_(ND4)_ | | - | - | TN93 | | Uniform [10^-10^ - 10^-6^] |
| Individual locus *µ*_(ND4)_ | | - | - | TN93 | | Gamma [10^-10^ - 10^-6^] |
| Mean *k1*_(ND4)_ | | - | - | TN93 | | Uniform [0.05 - 20] |
| Individual locus *k1*_(ND4)_ | | - | - | TN93 | | Gamma [0.05 - 20] |
| Mean *k2*_(ND4)_ | | - | - | TN93 | | Uniform [0.05 - 20] |
| Individual locus *k2*_(ND4)_ | | - | - | TN93 | | Gamma [0.05 - 20] |

N – effective population size for each analysed deme (EPY – eastern Pyrenees; CPY – central Pyrenees; CWPY – central-western Pyrenees; PEU – Picos de Europa Mountains, GUA – Guadarrama Mountains), ra – admixture rate, t – time of events in generations (t_1_ – time to the most recent split; t_2_ – time to the intermediate split; t_3_ – time to the most ancient split). Microsatellite (SSRs) and mitochondrial (ND4) parameters: mean *µ* – mean mutation rate, individual locus *µ* – individual locus mutation rate, mean *P* – mean coefficient *P*, individual locus *P* – individual locus coefficient *P*, SNI – Single Nucleotide Insertion rate, mean *k* – mean coefficient *k*, individual locus *k* – individual locus coefficient *k*. Microsatellite loci were divided in two groups depending on the motif length (tri- and tetranucleotide loci). Conditions: sequence data were simulated under a Tamura Nei (TN93) mutation model.
