## Supplementary material for "Contrasting patterns of genetic admixture explain the phylogeographic history of Iberian high mountain populations of midwife toads": S3 Table.docx

**S3 Table. Microsatellite-based (below diagonal) and ND4-based (above diagonal) pairwise estimates of F_ST_ between the seven genetic groups identified by STRUCTURE in *Alytes obstetricans*/*almogavarii*** **(see Fig 5).** All P values < 0.01.

|  | Eastern Pyrenees | Central Pyrenees | Central-western Pyrenees 1 | Central-western Pyrenees 2 | Picos de Europa | Guadarrama 1 – lowland | Guadarrama 2 |
| --- | --- | --- | --- | --- | --- | --- | --- |
| Eastern Pyrenees |  | 0.831 | 0.941 | 0.937 | 0.910 | 0.913 | 0.951 |
| Central Pyrenees | 0.098 |  | 0.744 | 0.642 | 0.681 | 0.710 | 0.746 |
| Central-western Pyrenees 1 | 0.122 | 0.097 |  | 0.302 | 0.727 | 0.820 | 0.906 |
| Central-western Pyrenees 2 | 0.176 | 0.192 | 0.093 |  | 0.688 | 0.781 | 0.914 |
| Picos de Europa | 0.092 | 0.107 | 0.095 | 0.155 |  | 0.776 | 0.858 |
| Guadarrama 1 – lowland | 0.106 | 0.132 | 0.102 | 0.151 | 0.060 |  | 0.089 |
| Guadarrama 2 | 0.175 | 0.195 | 0.157 | 0.222 | 0.115 | 0.084 |  |
