## Supplementary material for "Contrasting patterns of genetic admixture explain the phylogeographic history of Iberian high mountain populations of midwife toads": S4 Table.docx

**S4 Table. Analysis of molecular variance (AMOVA) for mitochondrial (ND4) and nuclear (microsatellites) markers based on the seven genetic groups identified by STRUCTURE in *Alytes obstetricans*/*almogavarii*** **(see Fig 5).** All P values < 0.001.

|  | ND4 |  |  | Microsatellites | |  |
| --- | --- | --- | --- | --- | --- | --- |
| Source of variation | SS | Variance component | % Variation | SS | Variance component | % Variation |
| Among clusters | 1149.248 | 6.259 | 84.945 | 1639.308 | 1.042 | 26.171 |
| Among populations within clusters | 148.165 | 0.466 | 6.319 | 1275.452 | 0.693 | 17.405 |
| Among individuals within populations | 77.883 | 0.644 | 8.736 | 3640.199 | 2.247 | 56.425 |
