## Supplementary material for "Contrasting patterns of genetic admixture explain the phylogeographic history of Iberian high mountain populations of midwife toads": S5 Table.docx

**S5 Table. Posterior probability of tested scenarios and 95% confidence intervals (CI) estimated with DIYABC analysis when considering only microsatellites and when including both mtDNA (ND4) and microsatellite markers.** Type I and II errors for the best supported scenarios are indicated. See Fig 2 for more information on tested scenarios.

|  | Microsatellites |  |  |  | Microsatellites + ND4 | | |  |
| --- | --- | --- | --- | --- | --- | --- | --- | --- |
| Scenario | Posterior probability | 95% CI | Type I error | Type II error | Posterior probability | 95% CI | Type I error | Type II error |
| 1 | 0.101 | 0.038-0.163 |  |  | 0.063 | 0.032-0.095 |  |  |
| 2 | 0.849 | 0.837-0.860 | 0.114 | 0.075 | 0.292 | 0.274-0.310 |  |  |
| 3 | 0.005 | 0.000-0.073 |  |  | 0.056 | 0.023-0.089 |  |  |
| 4 | 0.007 | 0.000-0.074 |  |  | 0.109 | 0.058-0.159 |  |  |
| 5 | 0.039 | 0.000-0.112 |  |  | 0.480 | 0.459-0.502 | 0.078 | 0.089 |
