## Supplementary material for "Contrasting patterns of genetic admixture explain the phylogeographic history of Iberian high mountain populations of midwife toads": S6 Table.docx

**S6 Table. Posterior parameters (median and 95% confidence intervals) and RMedAD (Relative Median Absolute Deviation) estimated with DIYABC analysis for the best supported scenarios when considering only microsatellites (simple sequence repeats – SSRs; scenario 2) and when including both mtDNA (ND4) and microsatellite markers (scenario 5).** See Fig 2 for more information on tested scenarios.

|  | Microsatellites | | | | Microsatellites + ND4 | | | |
| --- | --- | --- | --- | --- | --- | --- | --- | --- |
| Parameter | Median | *Q_2.5_* | *Q_97.5_* | RMedAD | Median | *Q_2.5_* | *Q_97.5_* | RMedAD |
| N_EPY_ | 12 100 | 6 110 | 18 500 | 0.159 | 15 800 | 10 500 | 19 200 | 0.201 |
| N_CPY_ | 8 830 | 3 820 | 17 300 | 0.195 | 18 200 | 14 300 | 19 800 | 0.202 |
| N_CWPY_ | 10 500 | 4 580 | 18 100 | 0.189 | 13 700 | 8 200 | 18 500 | 0.196 |
| N_PEU_ | 11 800 | 5 220 | 19 000 | 0.184 | 14 600 | 8 370 | 19 200 | 0.189 |
| N_GUA_ | 5 500 | 2 460 | 12 100 | 0.193 | 5 590 | 2 620 | 11 600 | 0.278 |
| N_EPY-GUA_ | 18 200 | 1 330 | 38 400 | 0.376 | - | - | - | - |
| N_EPY-CWPY-GUA_ | - | - | - | - | 11 700 | 742 | 35 900 | 0.486 |
| ra | - | - | - | - | 0.350 | 0.043 | 0.874 | 0.334 |
| t_1_ | 12 600 | 5 370 | 24 000 | 0.223 | 20 700 | 11 200 | 28 100 | 0.360 |
| t_2_ | 18 400 | 6 770 | 36 900 | 0.228 | 26 400 | 12 900 | 38 000 | 0.247 |
| t_3_ | 28 000 | 9 600 | 48 500 | 0.194 | 38 100 | 19 100 | 49 200 | 0.249 |
| Mean *µ1*_(SSRs)_ | 1.28x10^-4^ | 4.36x10^-5^ | 4.05x10^-4^ | 0.414 | 4.59x10^-5^ | 1.93x10^-5^ | 1.70x10^-4^ | 0.614 |
| Mean *P1*_(SSRs)_ | 0.216 | 0.111 | 0.294 | 0.258 | 0.177 | 0.104 | 0.287 | 0.222 |
| Mean *µ2*_(SSRs)_ | 3.89x10^-4^ | 1.86x10^-4^ | 8.07x10^-4^ | 0.328 | 3.31x10^-4^ | 1.49x10^-4^ | 7.41x10^-4^ | 0.536 |
| Mean *P2*_(SSRs)_ | 0.268 | 0.144 | 0.300 | 0.262 | 0.192 | 0.107 | 0.292 | 0.228 |
| Mean *µ*_(ND4)_ | - | - | - | - | 1.96x10^-7^ | 1.06x10^-7^ | 4.42x10^-7^ | 0.217 |
| Mean *k1*_(ND4)_ | - | - | - | - | 17.400 | 2.620 | 20.000 | 0.421 |
| Mean *k2*_(ND4)_ | - | - | - | - | 11.500 | 0.724 | 19.800 | 0.410 |

N – effective population size for each analysed deme (EPY – eastern Pyrenees; CPY – central Pyrenees; CWPY – central-western Pyrenees; PEU – Picos de Europa Mountains; GUA – Guadarrama Mountains), ra – admixture rate, t – time of events in generations (t_1_ – time to the most recent split; t_2_ – time to the intermediate split; t_3_ – time to the most ancient split), mean *µ* – mean mutation rate, mean *P* – mean coefficient *P*, mean *k* – mean coefficient *k,* *Q_2.5_* – quantile 2.5%, *Q_97.5_* – quantile 97.5%.
