## Supplementary figures and images for "Contrasting patterns of genetic admixture explain the phylogeographic history of Iberian high mountain populations of midwife toads"

### S2 Fig.pdf

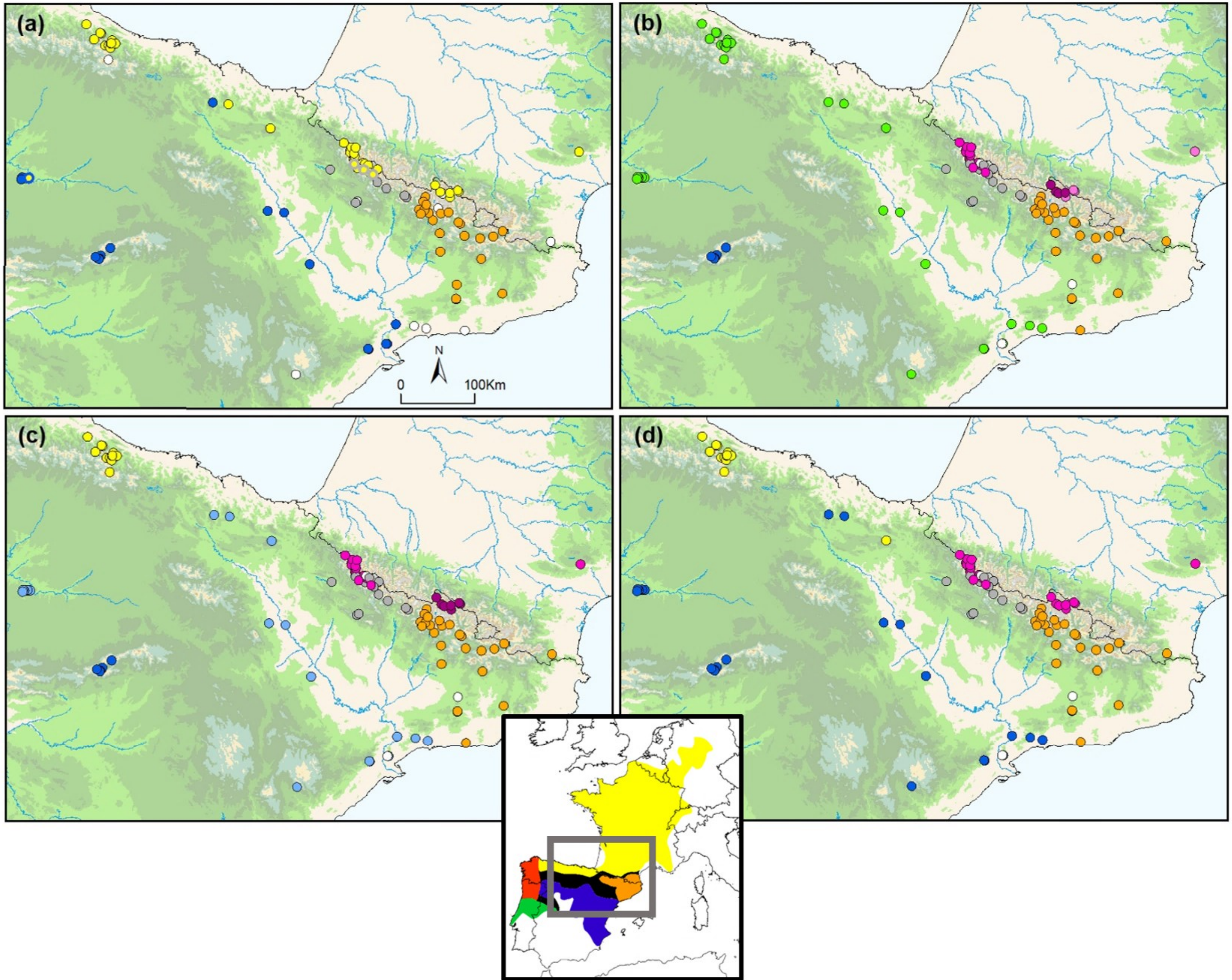

### S3 Fig.pdf

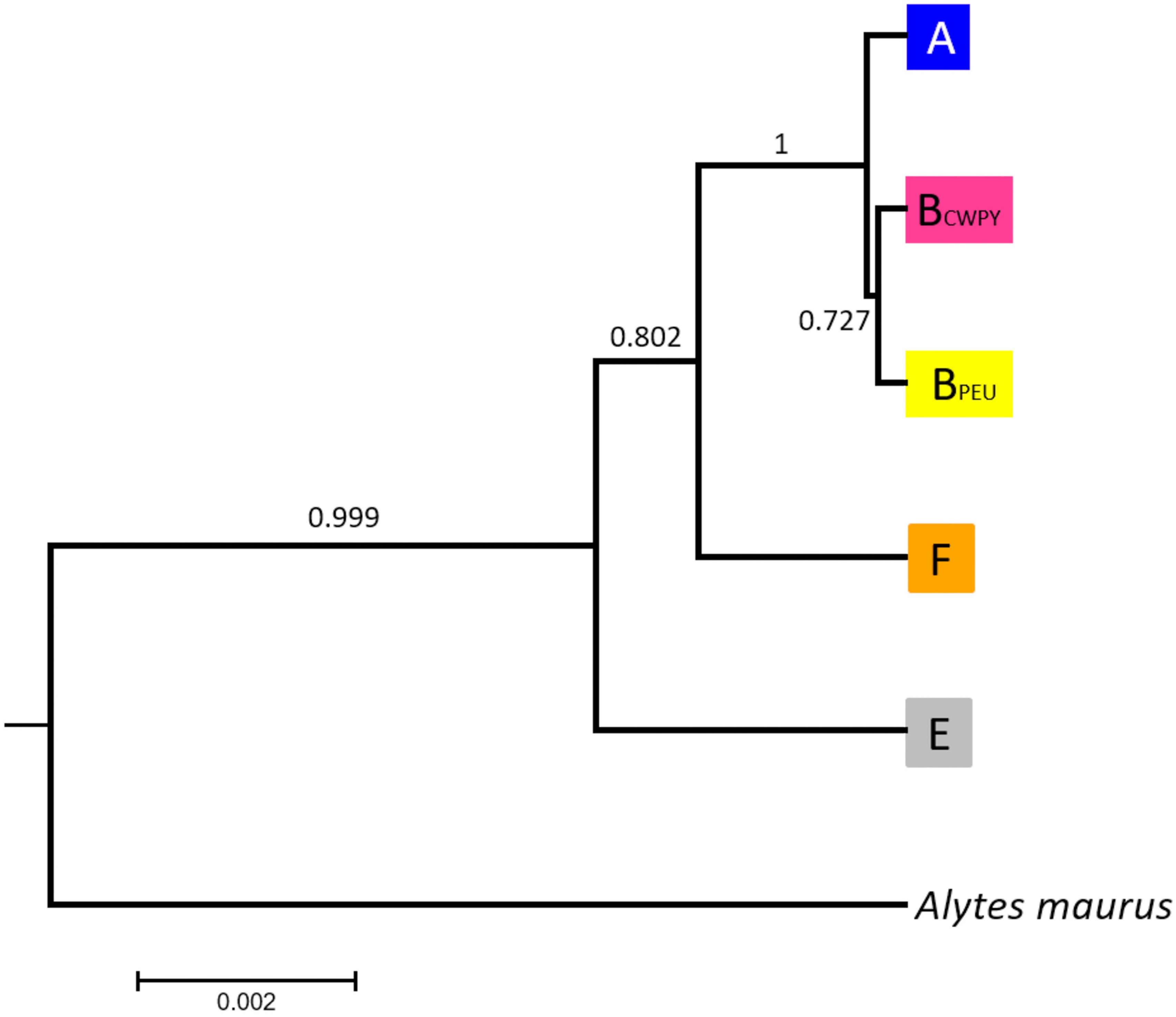

### S4 Fig.pdf

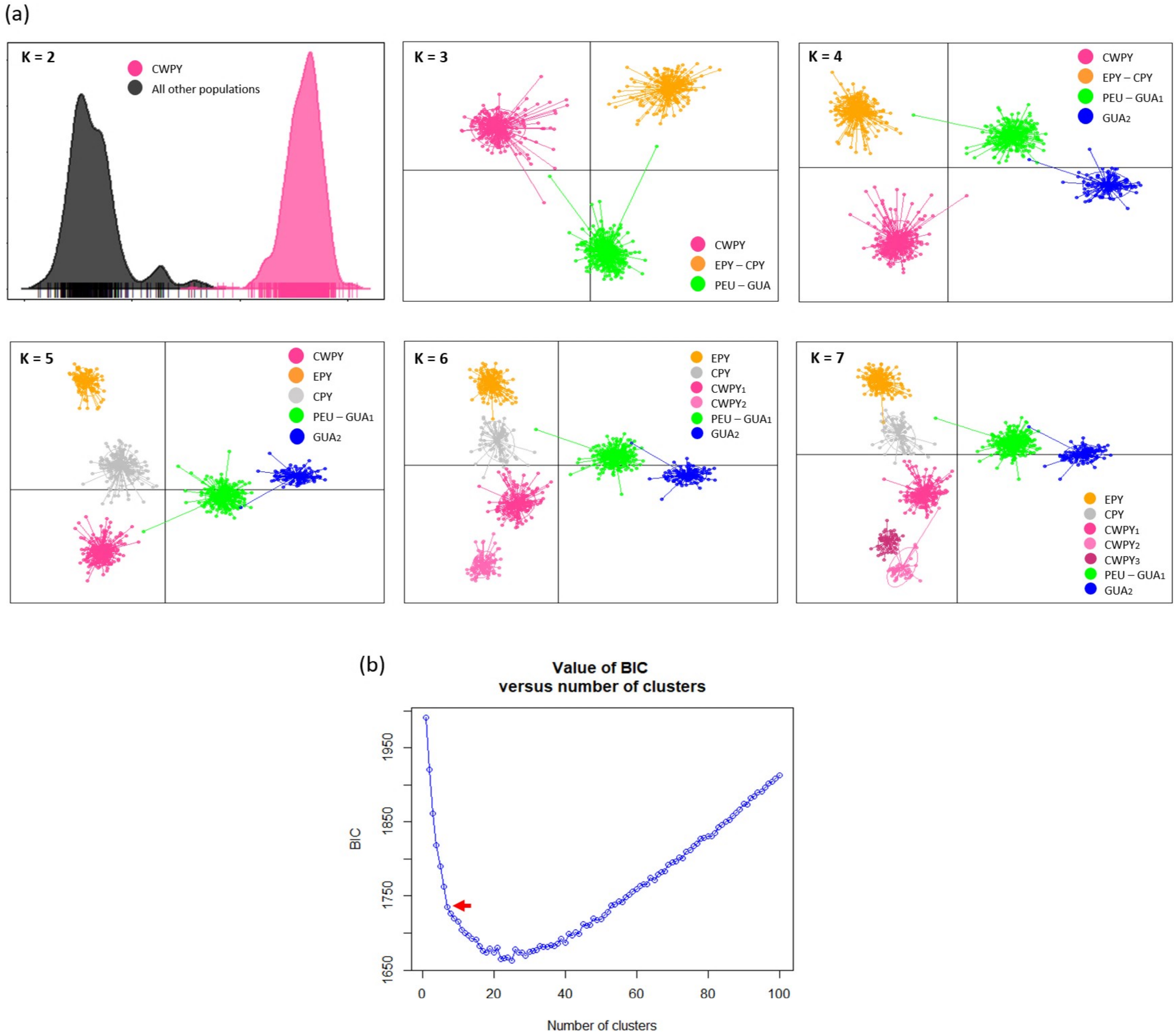

### S5 Fig.pdf

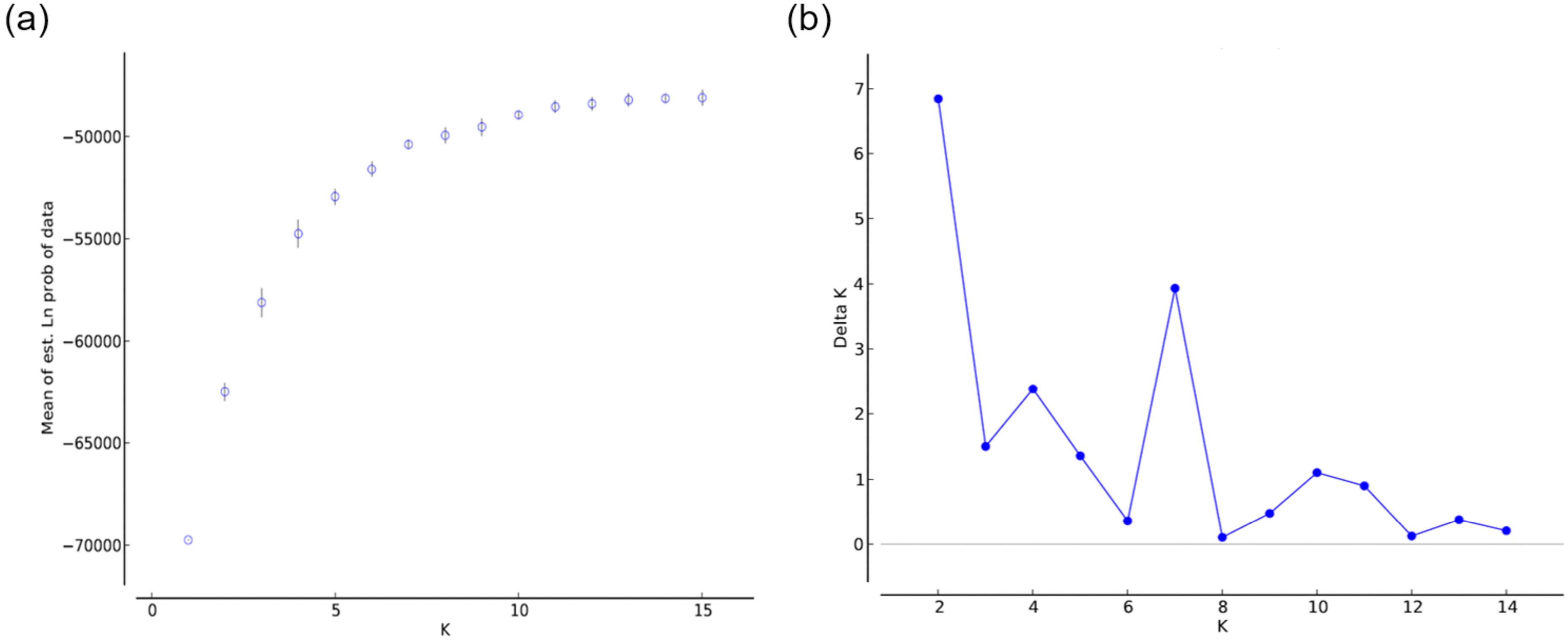

### S6 Fig.pdf

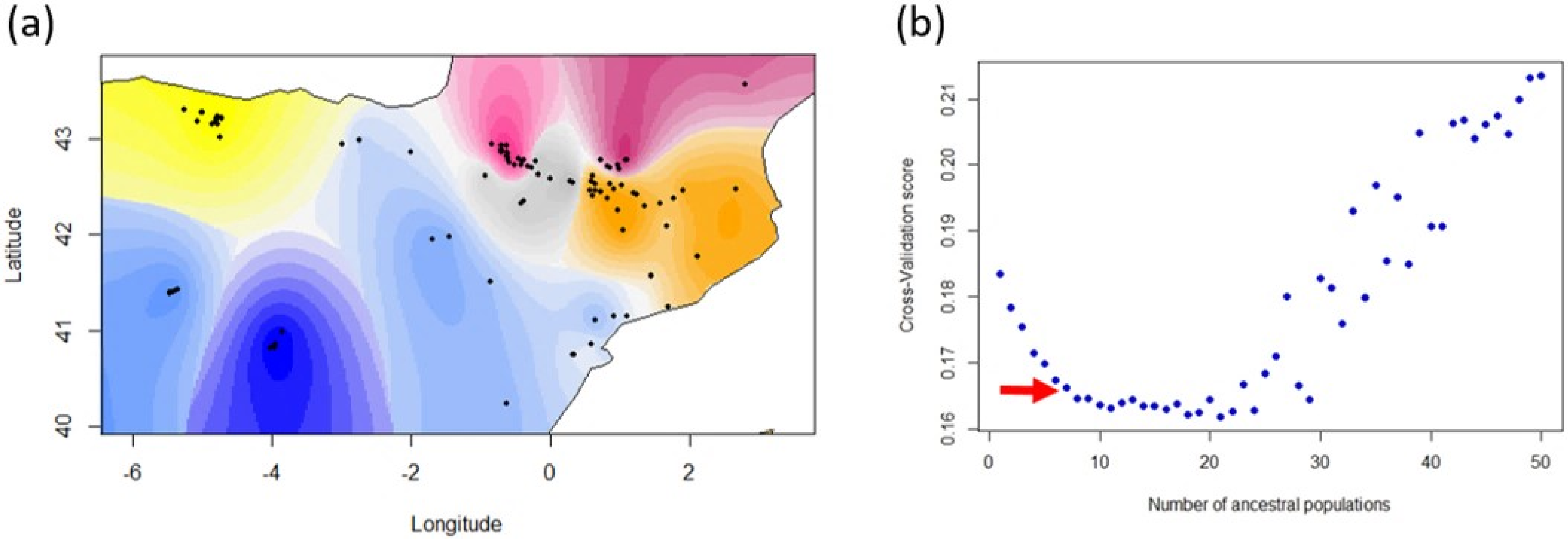

### S7 Fig.pdf

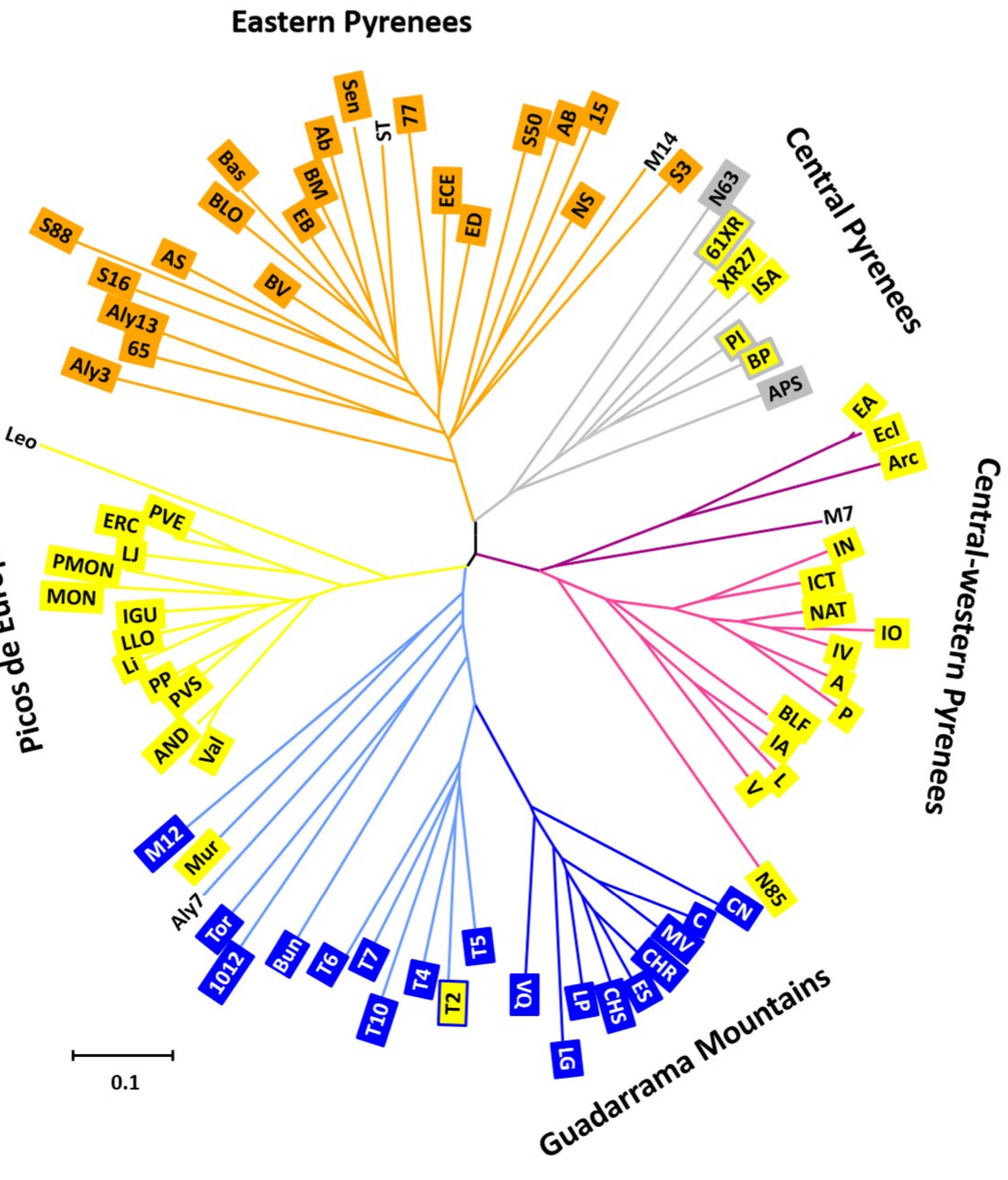
